# Spatiotemporal dynamics of DNA repair proteins between the Golgi and nucleus maintain genomic stability

**DOI:** 10.1101/2022.10.17.512236

**Authors:** George Galea, Karolina Kuodyte, Muzamil M. Khan, Peter Thul, Beate Neumann, Emma Lundberg, Rainer Pepperkok

## Abstract

The Golgi complex is a key organelle of the secretory pathway, but also serves as a critical hub for maintaining cellular homeostasis, orchestrating various signalling pathways and essential cellular processes such as membrane trafficking and post-translational modifications. Despite its central role, the communication between the Golgi and the cell nucleus has remained largely unexplored. To bridge this gap, we have analysed and siRNA-validated localisation data of the Human Protein Atlas, finding an unexpected and significant level of interconnectivity between the proteomes of the Golgi complex and nucleus, including an intriguing involvement in DNA repair processes. Here, we uncovered a cluster of DNA repair proteins present in distinct sub-Golgi localisations as well as in the nuclear compartment. In response to genotoxic stress these proteins redistribute dynamically between the Golgi complex and the nucleus, with the specific type of DNA injury influencing this distribution pattern. To probe this Golgi–nucleus link, we examined the Homologous Recombination (HR) regulator RAD51C and find that DNA damage triggers ATM-dependent release of a Giantin-tethered Golgi pool and Importin-β–dependent nuclear import, where repair-associated nuclear foci form. Downregulation of Giantin prematurely releases RAD51C, producing RAD51C nuclear foci lacking key HR markers, reducing ATM activation, elevating genome instability, and accelerating cell proliferation. These findings support a timing-based pathway in which the Golgi acts as a spatiotemporal coordination node for HR regulators and other DDR factors. Our study sheds light on the dynamic interplay between the Golgi and the nucleus in safeguarding genomic stability while maintaining the delicate balance of cellular homeostasis.

## Introduction

Eukaryotic cells have evolved a highly specialised and coordinated array of membrane-bounded organelles. This compartmentalisation allows the segregation of biochemical reactions, ensuring that they are carried out with the highest specificity and efficiency. However, organelles do not function in isolation but rely on the continual exchange of lipids, proteins and signalling cues to maintain cellular homeostasis. At the centre of this cross-coordination of subcellular transport and signalling pathways lies the Golgi complex. Thus, contributing well beyond its classical roles of membrane trafficking, and post-translational modification but also acting as a regulatory hub with numerous cellular processes intersecting at this organelle such as autophagy, mitosis, growth signalling, cytoskeletal and energy status regulation (Wilson et al. 2011; Makhoul, Gosavi, and Gleeson 2018). Perturbations to the Golgi architecture and mutations of its constituents have been associated with a wide array of human diseases such as neurodegenerative disorders and cancer, amongst many others (Freeze and Ng 2011; Machamer 2015; Zappa, Failli, and De Matteis 2018; Liu et al. 2021). Although we have a clearer picture of the Golgi’s interactions and regulatory functions in the cytoplasmic domain, its communication with the nucleus remains largely unexplored.

Emerging themes have started to illustrate the relationship between the Golgi and nuclear compartment, with multi-localisation proteins playing a pivotal role. For example, in cholesterol homeostasis and endoplasmic reticulum (ER) stress response, sensing-signalling proteins SREBP and ATF6, respectively, get proteolytically cleaved at the Golgi complex to regulate gene expression through the release of a transcriptionally active amino terminus that makes its way to the nucleus (Haze et al. 1999; Brown, Radhakrishnan, and Goldstein 2018). Few studies have also hinted at a link between cytoplasmic organelles, genomic stability, and, in turn, cancer, with the Golgi complex emerging as a central theme (Petrosyan 2015; Kulkarni-Gosavi, Makhoul, and Gleeson 2019; Zhang 2021). At a structural level, the Golgi undergoes dramatic morphological changes following the induction of DNA lesions, from ribbon-like perinuclear stack to dispersed fragments (Farber-Katz et al. 2014). A response requiring the phosphorylation of the Golgi resident oncoprotein, Golgi phosphoprotein 3 (GOLPH3), by DNA-dependent protein kinase (DNA-PK), a DNA Damage Response (DDR) regulator (Farber-Katz et al. 2014). Golgi morphology alterations are also a common feature across a wide variety of cancer types and are often reflected in changes in the distribution of Golgi resident proteins (Petrosyan 2015; Zhang 2021). Rearrangement of Golgi glycosyltransferases distribution is a recurrent phenomenon in cancer cells, resulting in defective glycosylation, a process thought to promote cancer development (Petrosyan 2015; Zhang 2021; Bui et al. 2021). These studies hint at a clear link between Golgi and nucleus as communicating organelles, however a systematic approach that could pin down the signalling pathways involved in this communication has been missing.

To this end, here, we utilise a localisation data and antibody resources from the Human Protein Atlas (HPA) project (Thul et al. 2017) to explore a class of multi-localisation proteins as a systematic strategy to identify key signalling pathways that function between the Golgi and nuclear compartment. We validate candidate Golgi–nuclear localisations by siRNA-mediated knockdown revealing a network of DNA repair proteins localised at the Golgi complex. We systematically analyse their subcellular localisation within the Golgi, as well as their redistribution patterns in response to different types of DNA damage. Building on these observations, we propose a spatiotemporal regulatory pathway for DDR control between the Golgi complex and nucleus that gates the timed availability of HR regulators. Using RAD51C as a representative HR factor, we find that a Golgi resident pool is released to the nucleus upon DNA damage where it forms repair associated foci, and that the Golgi scaffold Giantin (GOLGB1) is required for RAD51C localisation to the Golgi. Disrupting this regulation through Giantin depletion leads to premature nuclear accumulation of RAD51C, modulation of upstream ATM signalling, increased genomic instability, and accelerated cell proliferation. Together, these findings highlight the importance of spatial regulation of DDR proteins in maintaining genomic integrity and cellular homeostasis, and suggest a broader role for the Golgi as a regulatory platform in DNA repair.

## Results

### Antibody-based systematic analysis and siRNA-mediated validation identifies a network of DNA damage response proteins at the Golgi complex

To systematically explore candidates that might link Golgi to nuclear function or vice versa, we shortlisted 329 proteins annotated by the Human Protein Atlas (HPA) project (Thul et al. 2017) to localise at both Golgi-like membranes and the nucleus **(Figure 1A)**. To ensure the specificity of both localisation data and antibody binding, we tested the corresponding HPA antibody for each candidate using an siRNA-mediated knockdown pipeline (Stadler et al. 2012). This approach was specifically designed to exclude false localisation arising from antibody cross-reactivity. A reduction of at least 25% in both nuclear and Golgi signals following knockdown was required for validation.

**Figure 1:**
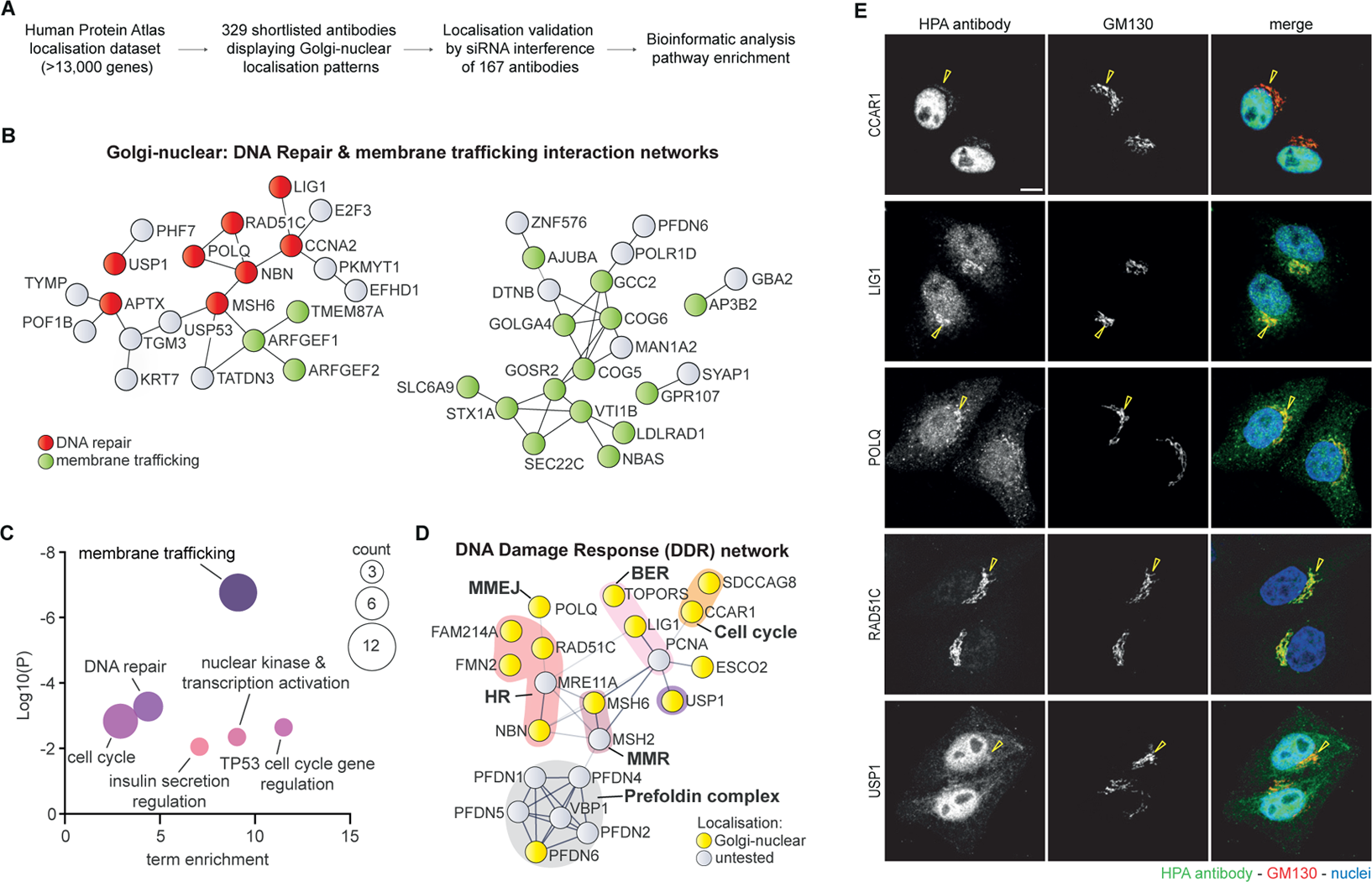
Antibody-based analysis identifies a network of DNA damage response (DDR) proteins at the Golgi complex. **(A)** Schematic diagram of the screening process for the shortlisting, validation and characterisation of Golgi-nuclear localisation proteins, as a strategy to identify linking pathways. **(B)** Experimentally-based protein-protein interaction network displaying the DNA repair and membrane trafficking Golgi-nuclear localisation proteins validated in this study; categorised by functional pathways. **(C)** Pathway enrichment analysis of the Golgi-nuclear localisation proteins identified in this study. **(D)** Experimentally-based STRING protein-protein interaction network showing the DDR proteins identified to localise to both the Golgi complex and nucleus; yellow nodes indicate double-localising proteins, and grey nodes are filler untested proteins. **(E)** Representative images of HeLa-K cells stained with Human Protein Atlas (HPA) antibodies against Golgi-nuclear DDR proteins (green), and the Golgi marker GM130 (red). DNA was stained with Hoechst 33342 (blue). Yellow arrowheads denote the Golgi complex. Scale bar, 10 μm.

Using this strategy, we confirmed the dual localisation for 163 proteins (167 HPA antibodies) **(Figure 1B and Table S1)**. Bioinformatic analysis of the identified 163 candidates revealed a number of functional protein networks **(Figure 1B)**, primarily composed of two major pathways: membrane trafficking and surprisingly, DNA Damage Response (DDR) **(Figures 1B and 1C)**. Additionally, we identified enrichment for cell cycle regulation, RNA metabolism, and lipid metabolism regulators **(Table S1 and Figure S2B)**. Within the membrane trafficking enriched cluster, we identified several essential components of intra-Golgi trafficking machinery, including Golgins, SNAREs, COG tethering complex, and RAB GTPase exchange factors **(Figure 1B)**.

An enriched cluster centred on the DDR emerged through the combination of DNA repair proteins and an array of cell cycle regulators (**Figures 1D, S1, S2A and S2B**). Within this cluster, we identified various proteins collectively spanning the core DNA repair pathways (Homologous Recombination (HR); Mismatch Repair (MMR); Microhomology-Mediated End Joining (MMEJ) and Base Excision Repair (BER)) as well as other integral regulators of DDR such as ubiquitination, cell cycle and signalling **(Figures 1D and 1E)**. These surprising findings of DNA repair proteins at the Golgi prompted us to explore the links between DDR and Golgi function.

### Golgi cisternal localisation analysis of dual-localising DDR proteins identifies specific distribution patterns that correlate with their function

To start addressing the meaning of DDR proteins localisation at the Golgi complex and further validate their localisation, we shortlisted 15 candidates based on their described role in this pathway and conducted an analysis of their subcellular distribution within the organelle. The Golgi apparatus consists of *cis*-, *medial*, and *trans*-Golgi cisternae, extending into a tubular *trans*-Golgi network (TGN), with resident proteins strategically distributed along the Golgi stack in order to serve their function (Kulkarni-Gosavi, Makhoul, and Gleeson 2019). Since the ribbon-like Golgi structures make subdomain mapping difficult, we used nocodazole treatment to disperse the ribbon into isolated mini-stacks, a standard approach that enables high-resolution analysis of cisternal distribution (Dejgaard et al. 2007). We then performed confocal microscopy with co-staining for the *cis*-Golgi marker GM130 and the *trans*-Golgi marker TGN46. Fluorescence intensity line profiles were acquired across individual mini-stacks, and the relative localisation of each DDR protein was quantified by calculating Pearson’s correlation coefficients (PCC) with the *cis*- and *trans*-markers (Dejgaard et al. 2007) **(Figures 2A–2E**, **S3D and Table S2**).

**Figure 2:**
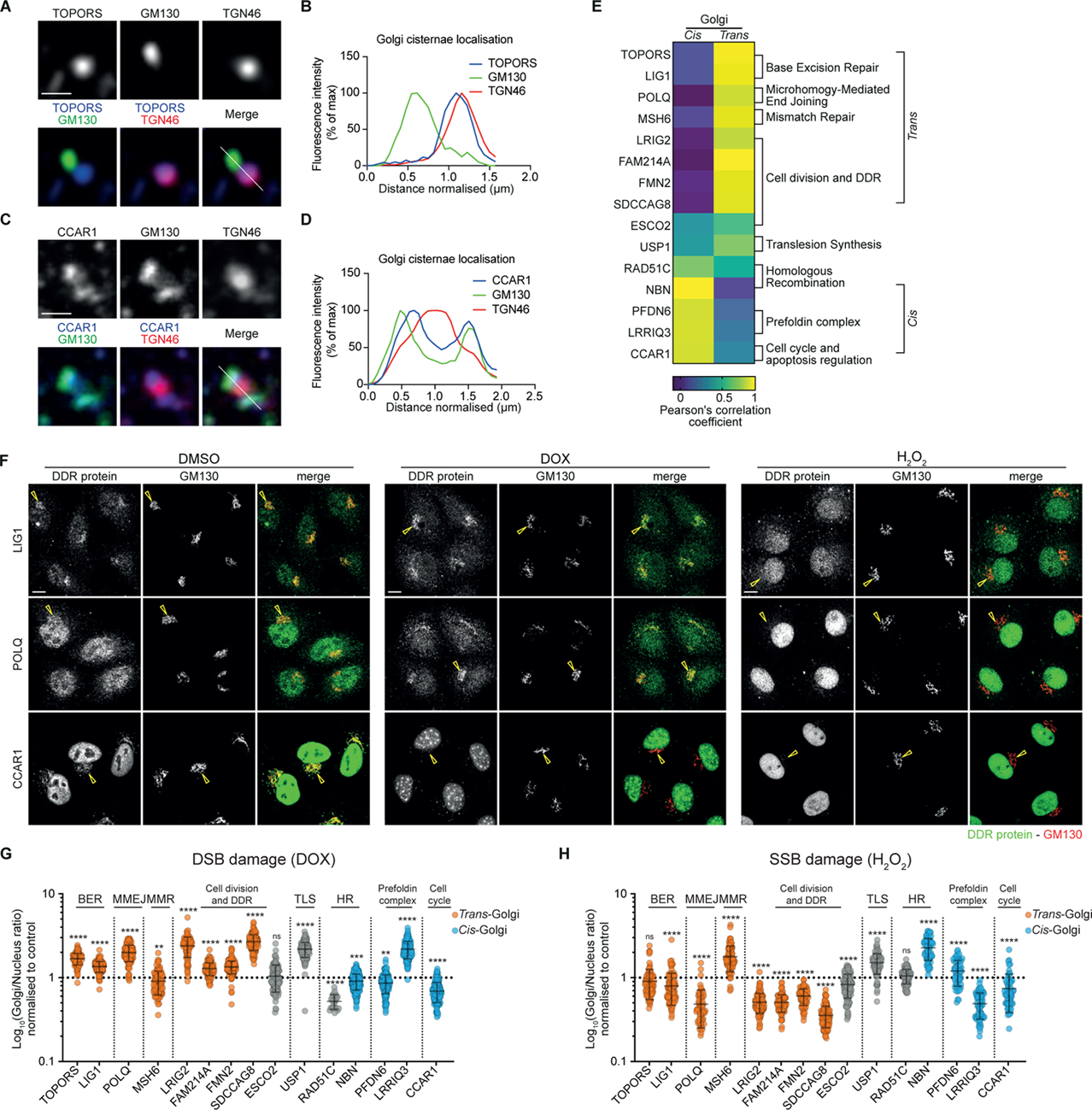
Systematic analysis of Golgi-nuclear DDR proteins localisation. **(A–E)** Golgi-cisternal localisation analysis of Golgi-nuclear DDR proteins. HeLa-K cells were treated with nocodazole (33 μM, 3 h), fixed and stained with antibodies against DDR proteins. **(A–C)** Enlarged view of single isolated mini-stack, where cells were stained with antibodies against TOPORS, CCAR1 and Golgi markers GM130 (*cis-*Golgi) and TGN46 (*trans-*Golgi); scale bar, 1 μm; **(B-D)** white lines across the stack depict the line-plot analysis. **(E)** Quantification of Pearson’s correlation coefficient (PCC) between *cis-*Golgi and *trans-*Golgi markers and DDR proteins; n ≥ 36 mini-stacks. **(F)** Localisation changes of dual-localising DDR proteins upon induction of DNA damage. HeLa-K cells were treated with the DNA damage-inducing drug, doxorubicin (DOX) (40 μM, 3 h) or oxidative stress-inducing agent hydrogen peroxide (H_2_O_2_) (50 μM for 20 min, followed with 15 min recovery), fixed and stained with antibodies against DDR proteins LIG1, POLQ, CCAR1 and the Golgi marker, GM130; yellow arrows denote the Golgi membranes. Scale bars, 10 μm. **(G)** A ratio of DDR protein Golgi-nuclear distribution after treatment with DOX. **(H)** A ratio of DDR protein Golgi-nuclear distribution after treatment with H_2_O_2_. Data represent the mean ± standard error of the mean (s.e.m.) (n = 3 biologically independent samples with at least 200 cells analysed for each protein in control and treatment conditions). The proteins are classified according to their Golgi localisation patterns **(E)**. Statistical significance was determined using a two-tailed unpaired Student’s t-test; ns, **P < 0.01, ***P < 0.001, ****P <0.0001, compared to untreated control.

This analysis resolved three major localisation patterns throughout the Golgi cisternae: (i) a subset preferentially correlating with the *trans*-Golgi marker, TGN46 **(Figures 2A, 2B and 2E**), (ii) proteins correlating with the *cis*-Golgi marker, GM130 **(Figures 2C, 2D and 2E**); and (iii) proteins without a strong preference for either marker. Although the number of proteins examined was limited, emerging trends suggest pathway-related distribution (**Figure 2E**). For instance, BER and MMEJ proteins TOPORS, LIG1, and POLQ exhibited co-localisation with the *trans-*Golgi marker, TGN46, while HR factors NBN (Nibrin) and RAD51C have a high correlation with the *cis-*Golgi marker, GM130.

These observations suggested that DDR proteins may not only localise to the Golgi but also occupy distinct cisternal subdomains, raising the possibility that sub-Golgi positioning could influence how different repair pathways respond to DNA damage. We therefore next examined whether these proteins dynamically redistribute between the Golgi and the nucleus in response to genotoxic stress.

### Dual-localising DDR proteins dynamically redistribute between the Golgi and nucleus in response to specific types of DNA injuries

We next examined whether DDR proteins with dual Golgi-nuclear localisation respond dynamically to DNA damage. To this end, we compared two classes of DNA damage: double-strand DNA breaks (DSBs) induced by doxorubicin (DOX), which intercalates into DNA (Thorn et al. 2011) and oxidative DNA lesions induced by hydrogen peroxide (H_2_O_2_) (Henle and Linn 1997) or potassium bromate (KBrO_3_) (Loft et al. 1998) **(Figures 2F–2H and S3– S5**).

Using this experimental approach we monitored for any genotoxic stress-induced changes in protein distribution pattern of the 15 Golgi-nuclear localising DDR proteins. Initial inspection revealed that all three treatments triggered marked changes in both the subcellular localisation distribution and the overall signal intensity of many of the 15 proteins tested. Two predominant redistribution patterns were observed: either in a Golgi-to-nuclear manner or conversely, in a nucleus-to-Golgi direction. To quantify these shifts, we measured fluorescence intensities in Golgi and nuclear masks and calculated a Golgi-to-nuclear distribution ratio for each protein. Ratios <1 indicate Golgi-to-nuclear redistribution, whereas ratios >1 reflect nucleus-to-Golgi shifts. This approach normalised for differences in protein abundance and cell-to-cell heterogeneity.

The quantifications of the Golgi-nuclear ratio revealed a correlation between the redistribution patterns of these DDR proteins and their subcellular distribution with the Golgi complex. Here, treatment with DOX **(Figures 2F, 2G and S3**) resulted in a predominantly uniform localisation pattern change, where *cis*-Golgi DDR proteins (3 out of 4) shifted from the Golgi to the nucleus, whereas *trans*-Golgi proteins (7 out of 8) displayed a redistribution pattern from the nucleus to the Golgi. Conversely, exposure to H_2_O_2_ or KBrO_3_ **(Figures 2F, 2H, S4 and S5**) elicited the opposite response, with the majority of *trans*-Golgi DDR proteins (7 out of 8) displaying a redistribution from the nuclear compartment to the Golgi while 2 out of the 4 *cis*-Golgi DDR proteins displayed a redistribution pattern from the nucleus to the Golgi.

When analysed by pathway annotation, broad distinctions also emerge, BER proteins (TOPORS and LIG1) and MMEJ factor POLQ shifted from nucleus to Golgi in response to DOX, whereas the MMR protein MSH6 and HR-associated proteins (RAD51C and NBN) redistributed from Golgi to nucleus **(Figures 2G and S3)**. Specifically, with DOX treatment, BER and MMEJ proteins shifted from the nucleus to the Golgi, while MMR and HR proteins relocated from the Golgi to the nucleus. Conversely, this trend was reversed with H_2_O_2_ or KBrO_3_ treatments **(Figures 2H, S4 and S5)**.

While pathway-level trends were evident, some proteins (for example CCAR1) exhibited similar directionality under DOX and H₂O₂, underscoring that pathway membership is not exclusive and many DDR factors act in multiple contexts. Several proteins participate in multiple repair contexts, complicating binary distinction between different pathways. Nonetheless, these findings demonstrate that DDR proteins redistribute between Golgi and nucleus in a manner that depends on both the type of DNA lesion and their steady-state Golgi cisternal localisation.

### Redistribution of RAD51C Golgi fraction is required for the formation of RAD51C repair nuclear foci and is dependent on the kinase ATM

To mechanistically dissect the novel link between the Golgi complex and the nucleus, we characterised the Homologous Recombination (HR) repair protein RAD51C. As a member of the RAD51 paralog family, RAD51C is essential for regulating HR-mediated repair of double-strand breaks (DSBs). Previous studies have demonstrated RAD51C’s involvement across multiple stages of HR repair: promoting the DNA damage checkpoint (Badie et al. 2009), aiding in RAD51 filament formation, stabilising replication forks, and resolving Holliday junctions to complete repair (Prakash et al. 2022; Rawal et al. 2023; Greenhough et al. 2023). RAD51C also functions in replication stress responses (Somyajit, Subramanya, and Nagaraju 2012) underscoring its versatility. This multifunctionality, combined with a substantial Golgi-localised population compared to other DDR proteins in our screen, prompted us to investigate RAD51C as a representative candidate for understanding how Golgi-associated DDR factors are dynamically recruited to repair complexes. Notably, RAD51C contains both a nuclear localisation signal (NLS) and a nuclear export signal (NES) (Miller et al. 2005), which may enable it to shuttle between the Golgi and the nucleus in response to DNA damage, thereby supporting its role in replication fork stabilisation and HR repair.

Following the identification of RAD51C as a dual-localising DDR protein in our initial HPA-based antibody screen, we sought to further validate its localisation using two additional commercial antibodies targeting distinct RAD51C epitopes. In total, three distinct antibodies were used: the original HPA antibody from the primary screen, a second antibody for independent immunofluorescence validation, and a third antibody for biochemical fractionation. Our immunofluorescence across across several cell lines confirmed RAD51C enrichment in a juxtanuclear compartment co-localising with the Golgi marker GM130 (**indicated by the yellow arrowhead; Figures 3A and 3B**), together with diffused cytoplasmic staining and discrete nuclear foci (**indicated by white arrowheads; Figures 3A and 3B)**. The specificity of the antibodies was confirmed by the depletion of RAD51C in HeLa Kyoto (HeLa-K) cells (**Figures S6A–S6C**). To corroborate these results biochemically, we fractionated HeLa-K cell lysates into nuclear, membrane and cytoplasmic compartments. RAD51C was detected in all fractions, with the majority present in the membrane and cytoplasmic pool (**Figures 3C and 3D**) consistent with the immunofluorescence data.

**Figure 3:**
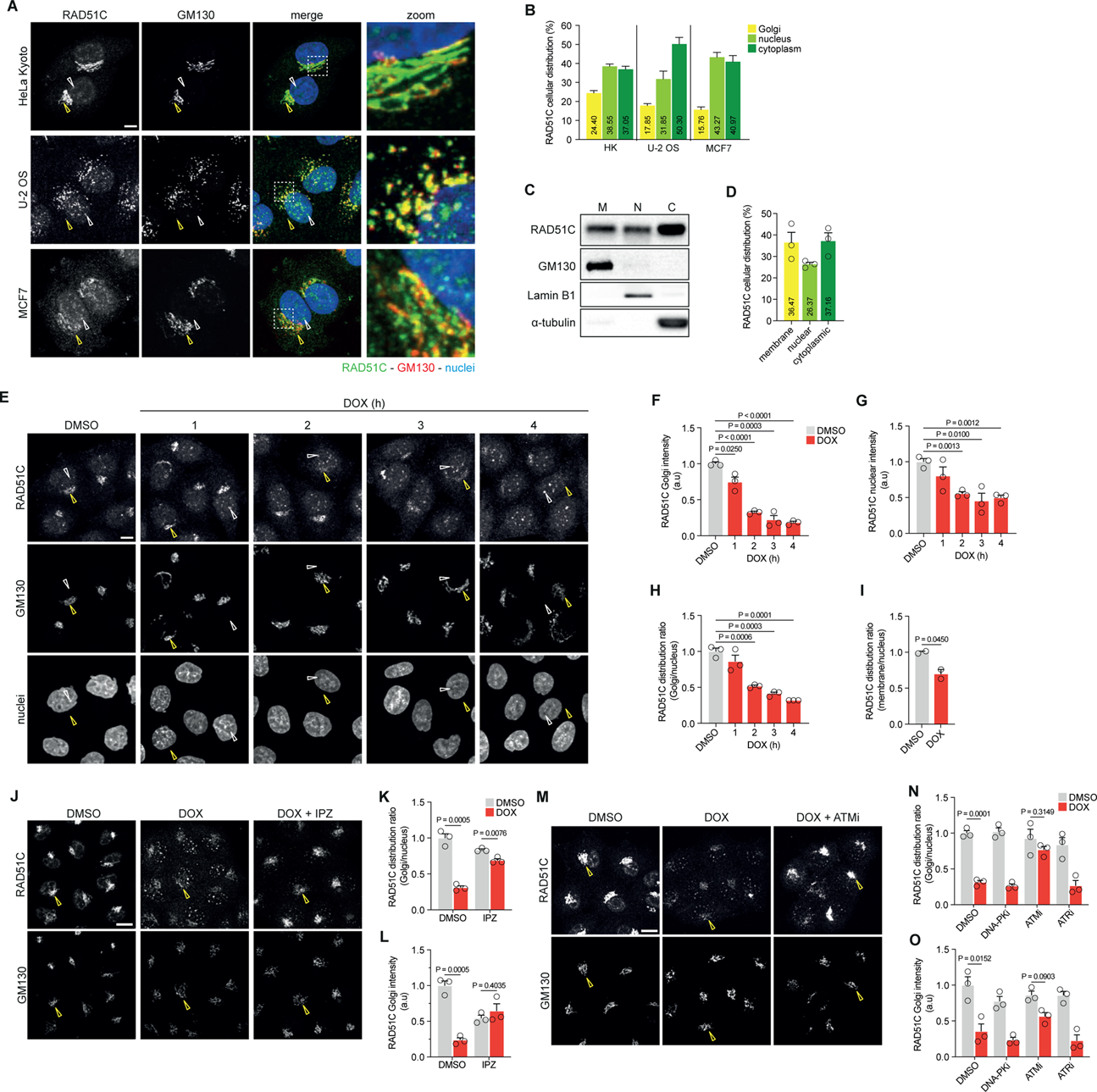
Redistribution of RAD51C Golgi fraction is required for the formation of RAD51C nuclear foci and is dependent on the kinase ATM. **(A)** Representative images of HeLa-K, U-2 OS and MCF7 cells stained with antibodies against RAD51C (green) and GM130 (red). DNA stained with Hoechst 33342 (blue). **(B)** Quantification of RAD51C distribution across the Golgi, nucleus and cytoplasm in various cell lines quantified from immunofluorescence images. **(C)** WB analysis showing the subcellular membrane (M), nuclear (N) and cytoplasmic (C) fractions of RAD51C (markers: GM130 for Golgi membranes, Lamin B1 for the nuclear compartment, alpha-tubulin for the cytoplasmic fraction). **(D)** Quantification of western blot analysis showing the distribution of RAD51C in 3 cell fractions: membrane, nuclear and cytoplasmic; (n = 3 biologically independent experiments) data represent the mean ± s.e.m. **(E)** HeLa-K cells treated with the DNA damage-inducing drug, doxorubicin (DOX), for increasing lengths of time. Yellow arrows denote the Golgi membrane; white arrows denote nuclear foci. **(F)** Quantification of sum intensity of RAD51C Golgi population, **(G)** RAD51C nuclear population, and **(H)** a ratio of RAD51C Golgi-nuclear distribution after treatment with DOX. Data represent the s.e.m (n = 3 biologically independent samples with a total of 2343 cells analysed). **(I)** Quantification of RAD51C membrane-nuclear distribution ratio after treatment with DOX, calculated from isolated fractions **(Figure S6E)**. HeLa-K cells were stained with antibodies against RAD51C and GM130. **(J)** Cells were treated with DMSO or IPZ prior to a 3-hour treatment with DOX. Yellow arrows denote the Golgi membrane; white denote nuclear foci. Results were quantified as **(K)** ratio of RAD51C Golgi-nuclear distribution and **(L)** relative sum intensity of RAD51C at the Golgi Data represents s.e.m. (n = 3 biologically independent samples with a total of 445 cells analysed). **(M)** HeLa-K cells were treated with DMSO (control) alone or with ATM inhibitor (KU55933) or ATR inhibitor (VE-821) or DNA-PK inhibitor (NU7441) prior to a 3-hour treatment with doxorubicin. Results were quantified as **(N) r**atio of RAD51C Golgi-nuclear distribution and **(O)** relative sum intensity of RAD51C at the Golgi. Data represents s.e.m. (n = 3 biologically independent samples with a total of 1679 cells analysed). Scale bars, 10 μm. Statistical significance was determined using a two-tailed unpaired Student’s t-test.

Although RAD51C has been extensively characterised biochemically and structurally, previous studies have examined its subcellular distribution. Earlier work has largely focused on *in vitro* complex formation and DNA repair mechanisms, rather than on the spatial distribution, likely contributing to the Golgi-associated pool remaining unrecognised. Although previous work, notably *Badie et al.* (2009), has shown a juxtanuclear RAD51C signal (see Fig. 1D of that study), this localisation was not further discussed.

Having established the steady-state RAD51C Golgi localisation, we next examined RAD51C dynamics following DSB induction with doxorubicin (DOX). Upon drug addition (**Figures 3E-3H**), we observed an overall decrease in RAD51C protein level, with the Golgi-localised fraction (co-localised with GM130, **yellow arrows**) decreasing more rapidly than the nuclear fraction. Concurrently, the diffused nuclear RAD51C pattern changed into very distinct nuclear foci (**white arrows**), which become more pronounced with longer DOX exposure (**Figure 3E**).

Quantitative analysis of these experiments allowed us to measure the changes in RAD51C levels at both the Golgi (**Figure 3F**) and the nuclear compartment (**Figure 3G**), and to calculate a RAD51C Golgi-nuclear distribution ratio (**Figure 3H**). This redistribution was further validated by subcellular fractionation (**Figures 3I and S6D**). Similarly, the total protein level of RAD51C is observed to decrease after 3 h treatment, with a much larger reduction in the membrane compartment when compared to the nuclear fraction. Additionally, we observed an increase in RAD51C cytoplasmic fraction with DOX treatment. Overall, in both experiments, the RAD51C Golgi-nucleus distribution ratio is seen to decrease significantly after 3 h DOX treatment in both the immunofluorescence and biochemical assay (**Figures 3H and 3I)**.

To identify a mechanism for RAD51C redistribution between compartments, we induced DSBs using DOX while inhibiting Importin-β-mediated nuclear import with the small peptide inhibitor Importazole (IPZ) (Soderholm et al. 2011). IPZ treatment (**Figures 3J–3L and S7A; Golgi fraction marked with yellow arrows; nuclear foci marker with white arrows**) inhibited the DOX-induced RAD51C redistribution. Instead, the majority of the protein population remained co-localised with the Golgi marker, GM130, and nuclear foci formation was significantly inhibited **(Figure 3J)**. While a reduction in the overall RAD51C protein level was measured with DOX treatment alone, no significant change in the RAD51C distribution pattern was observed with IPZ treatment (**Figure S7A)**. Unexpectedly, although DOX alone triggers dissociation of the Golgi RAD51C pool, under DOX plus IPZ RAD51C remained Golgi localised, suggesting that Importin-β dependent transport may contribute to or be tightly coupled with the RAD51C dissociation step.

We next sought to test whether the phosphorylation of DDR kinases mediate RAD51C redistribution in response to DSBs. The three master kinases that are active in response to DNA damage are ataxia telangiectasia mutated (ATM), ataxia telangiectasia and Rad3- related (ATR) and DNA-dependent protein kinase (DNA-PK) (Ciccia and Elledge 2010). Cells were treated with phosphorylation inhibitors specific for each kinase and RAD51C redistribution was analysed following DOX treatment (**Figures 3M–3O and S7B; Golgi fraction marked with yellow arrows**). DOX combined with the ATM phosphorylation inhibitor, KU55933 (Hickson et al. 2004) significantly inhibited RAD51C redistribution, with the majority of the protein remaining co-localised with GM130 and nuclear foci formation blocked **(Figure 3M)**. In contrast, the combined treatment of DOX with ATR or DNA-PK phosphorylation inhibitors, VE-821 (Fokas et al. 2012) and NU7441 (Tavecchio et al. 2012) respectively, had no apparent impact on the redistribution of RAD51C when compared with cells treated with DOX only. Inhibitor-only treatments did not alter RAD51C localisation (**Figure S7B**).

To confirm that these observations were a direct result of DSBs and not an off-target drug effect, we tested other DSB-causing agents (Jekimovs et al. 2014): camptothecin (CPT) (**Figures S8A–S8C**), etoposide (ETO) (**Figures S8D–S8F**), and mitomycin C (MMC) (**Figures S8G–S8I**). All treatments led to significant RAD51C redistribution from the Golgi to the nuclear compartment. Notably, CPT treatment (**Figure S8A**) caused a shift in nuclear RAD51C from a **diffuse distribution to distinct foci**, while ETO or MMC treatments (**Figures S8D and S8G**) increased the nuclear RAD51C population without obvious foci formation. These distinct nuclear phenotypes likely reflect lesion class and timing: CPT generates replication associated breaks that favour discrete HR foci, whereas ETO (Topo II poisoning) and MMC (interstrand crosslinks) produce damage that, within our treatment window, increases nuclear RAD51C without prominent puncta.

### Golgi localisation of RAD51C is dependent on the Golgin Giantin

To identify potential Golgi membrane anchors for RAD51C, we re-analysed two independent, previously published genome-wide siRNA datasets that profiled HR and DDR regulators (Paulsen 2010; Adamson et al. 2012). Interestingly these screens contained multiple Golgin protein family members as hit candidate genes, in particular the knockdown of Giantin and GMAP210 led to a significant reduction in HR repair rates (Adamson et al. 2012) (**Golgin results summarised in Figure S9A**), and the depletion of Giantin significantly inhibited H2AX phosphorylation, a crucial event in DDR signalling regulation (Paulsen 2010) (**Golgin results summarised Figure S9B**). Given the structural properties of Golgins (**Figure 4A**), which are predominantly coiled-coil proteins anchored to the Golgi membrane by their carboxy terminus and projected into the surrounding cytoplasm (Witkos and Lowe 2015), they are ideally suited for capturing or tethering nearby membranes and potentially retaining DDR proteins such as RAD51C.

**Figure 4:**
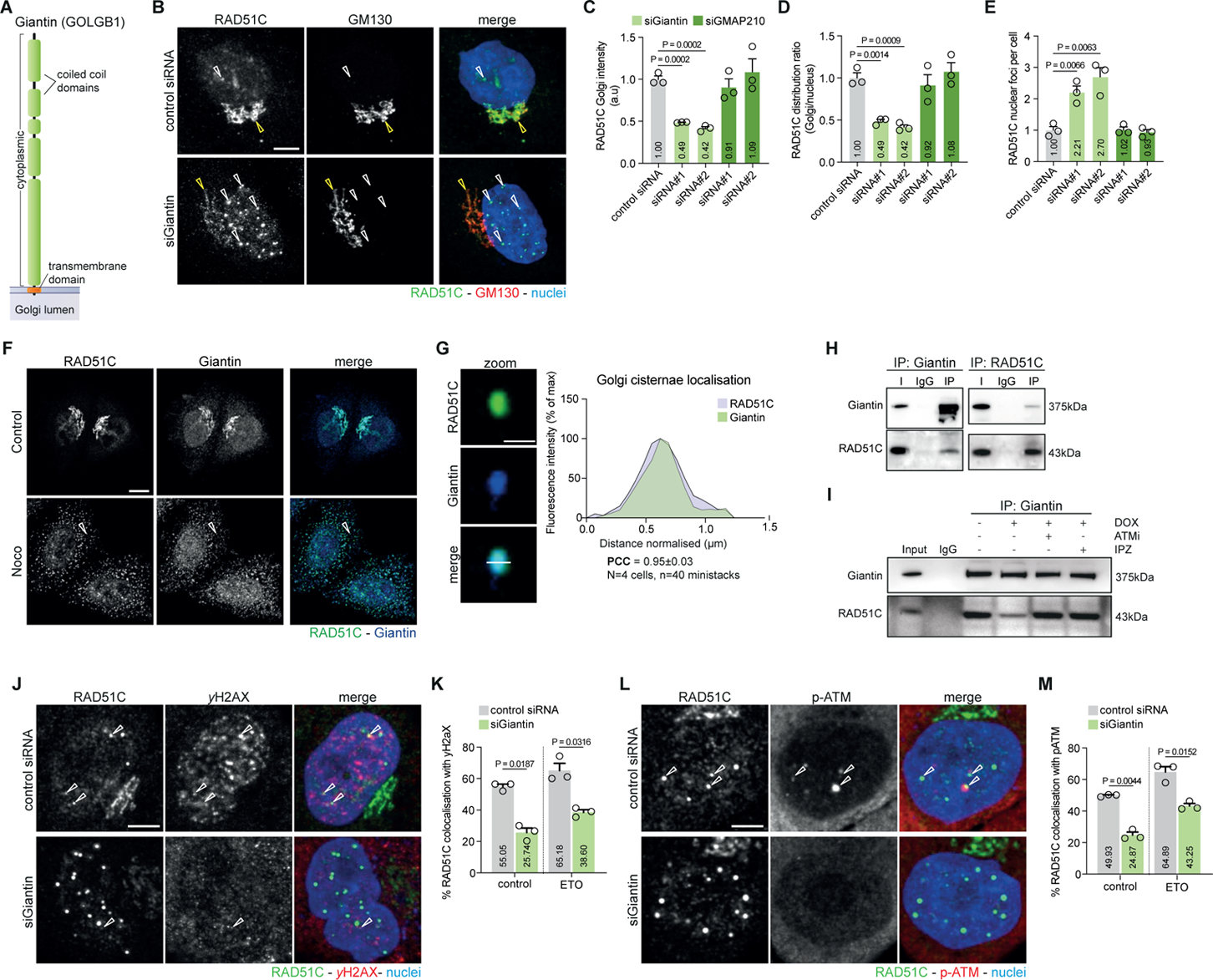
RAD51C Golgi localisation is dependent on the Golgin protein, Giantin. **(A)** Schematic diagram highlighting the domain organisation of Giantin. **(B)** RAD51C protein redistribution upon depletion of Giantin. HeLa-K cells were transfected with either control or Giantin siRNAs for 72 h, then immunostained for RAD51C and the Golgi marker GM130. Yellow arrows mark the Golgi, and white arrows denote RAD51C nuclear foci. **(C–E)** Quantifications of RAD51C localisation: **(C)** total Golgi-associated RAD51C intensity, **(D)** the Golgi-to-nucleus RAD51C intensity ratio, and **(E)** the number of nuclear RAD51C foci per cell. Data are presented as mean ± s.e.m. (n = 3 biologically independent experiments, 1,445 total cells). **(F)** Golgi-cisternal co-localisation analysis. HeLa-K cells treated with Nocodazole (33 μM, 3 h), fixed and stained with antibodies against Giantin and RAD51C. **(G)** Zoom enlarged view of single isolated mini-stack; scale bar, 1 μm; white line across the stack was used for line-scan analysis. **(H and I)** Co-immunoprecipitation (co-IP) of endogenous Giantin with RAD51C under control conditions **(H)** or after doxorubicin (DOX) treatment and co-treatments with ATM inhibitor (ATMi) and importazole (IPZ) **(I)** in HeLa-K extracts. **(J–M)** Co-localisation experiment of cells treated with control siRNA, or Giantin siRNA under control conditional or after treatment with ETO. Cells stained with antibodies against RAD51C (green) and HR DDR markers (red): **(J)** γ-H2AX, and **(L)** p-ATM. Co-localisation of structure is denoted by an arrowhead. Quantification of percentage RAD51C foci co-localising with **(K)** γ-H2AX and **(M)** p-ATM. Data represent the mean ± s.e.m. (n = 3 biologically independent samples with at least 200 cells and at least 550 RAD51C nuclear foci analysed for the co-localisation experiments for each condition).

To investigate whether RAD51C localisation is dependent on these Golgins, we performed siRNA-mediated depletion of Giantin and GMAP210, and compared their effects on RAD51C distribution to a control siRNA treatment (**Figures 4B–4E**). GMAP210 depletion had no significant effect on RAD51C distribution, whereas Giantin knockdown led to a notable redistribution of RAD51C (**Figure 4B**). Specifically, we observed a marked decrease in RAD51C co-localisation with the Golgi marker GM130 (**yellow arrows**) and an increase in nuclear RAD51C localisation with distinct bright nuclear foci appearing (**white arrows**) (**Figure 4B**). Quantifications revealed a significant reduction of the RAD51C Golgi population, (**Figure 4C**) and the distribution ratio (**Figure 4D**) reduced by more than half. The number of RAD51C foci (**Figure 4E**) increased by more than two-fold upon depletion of Giantin with either siRNAs. There was no significant difference in either RAD51C Golgi intensity, distribution ratio or foci number (**Figures 4C–4E**) between the GMAP210-depleted and control cells. The effective knockdown of Giantin (**Figure S9C**) and GMAP210 was tested by immunofluorescence. Sub-Golgi cisternae localisation analysis confirmed that Giantin and RAD51C are distributed in a similar manner throughout the organelle (**Figure 4F and G**).

To determine if RAD51C physically interacts or is part of a larger complex with Giantin we performed an immunoprecipitation (IP) assay (**Figures 4H and 4I**). Endogenous Giantin and RAD51C were found to co-IP from HeLa-K protein extracts (**Figure 4H**). To explore the dynamics of this interaction in response to DNA damage, we induced DSBs and performed co-IP experiments (**Figure 4I**). Our results showed that RAD51C and Giantin dissociate upon DSB induction. However, co-treatment with either an ATM inhibitor or IPZ, alongside doxorubicin, prevented this dissociation, corroborating our earlier immunofluorescence observations (**Figures 3E–3O**).

To gain insight into the nature of RAD51C foci induced by Giantin depletion, we carried out co-localisation assays to determine whether these structures contain standard HR markers, phosphorylated H2AX (γ-H2AX) (**Figures 4J and 4K**) and phosphorylated ATM (p-ATM) (**Figures 4L and 4M**) under physiological conditions and induction of DSBs by ETO. Both proteins are well-established markers for DSB repair sites and are important for the recruitment of the HR repair machinery (Vítor et al. 2020). In cells treated with a control siRNA, we observed that approximately half of the RAD51C nuclear foci were decorated with either γ-H2AX or p-ATM (**Figures 4J–4M**; **colocalising foci are denoted with an arrow**), as previously described (Vítor et al. 2020). RAD51C foci induced by the depletion of Giantin, however, showed significantly lower co-localisation with both markers regardless of DSBs induction. Overall, Giantin-depleted cells showed fewer γ-H2AX and p-ATM nuclear foci, consistent with previous reports (Paulsen 2010) (**Figure S9B**).

### Giantin depletion increases genomic instability and cell proliferation while inhibiting ATM signalling

Having built evidence that Giantin regulates Golgi tethering of RAD51C and its DNA damage–triggered release, we asked whether downregulation of Giantin impacts genome stability and DDR signalling. We analysed GMAP210 alongside Giantin because prior HR/DDRs screens implicated both Golgins (Paulsen 2010; Adamson et al. 2012). As a first readout, we assessed micronuclei formation, a well-established marker of genotoxic stress (**Figures 5A–5C**) (Krupina, Goginashvili, and Cleveland 2021). Under basal conditions, depletion of Giantin or RAD51C significantly increased the percentage of cells displaying micronuclei and aberrant nuclear structures compared to controls, while GMAP210 knockdown had no significant effect (**Figure 5B**). To test repair capacity, we induced DNA damage and allowed recovery (**Figure 5C**). Strikingly, after recovery both GMAP210 and Giantin knockdown led to a sustained and significant increase in micronuclei formation post-recovery, indicating persistent genomic instability and defective repair dynamics. Because our mechanistic model centres on a Giantin–RAD51C tether and the strongest basal phenotype mapped to Giantin, subsequent assays focus on Giantin.

**Figure 5:**
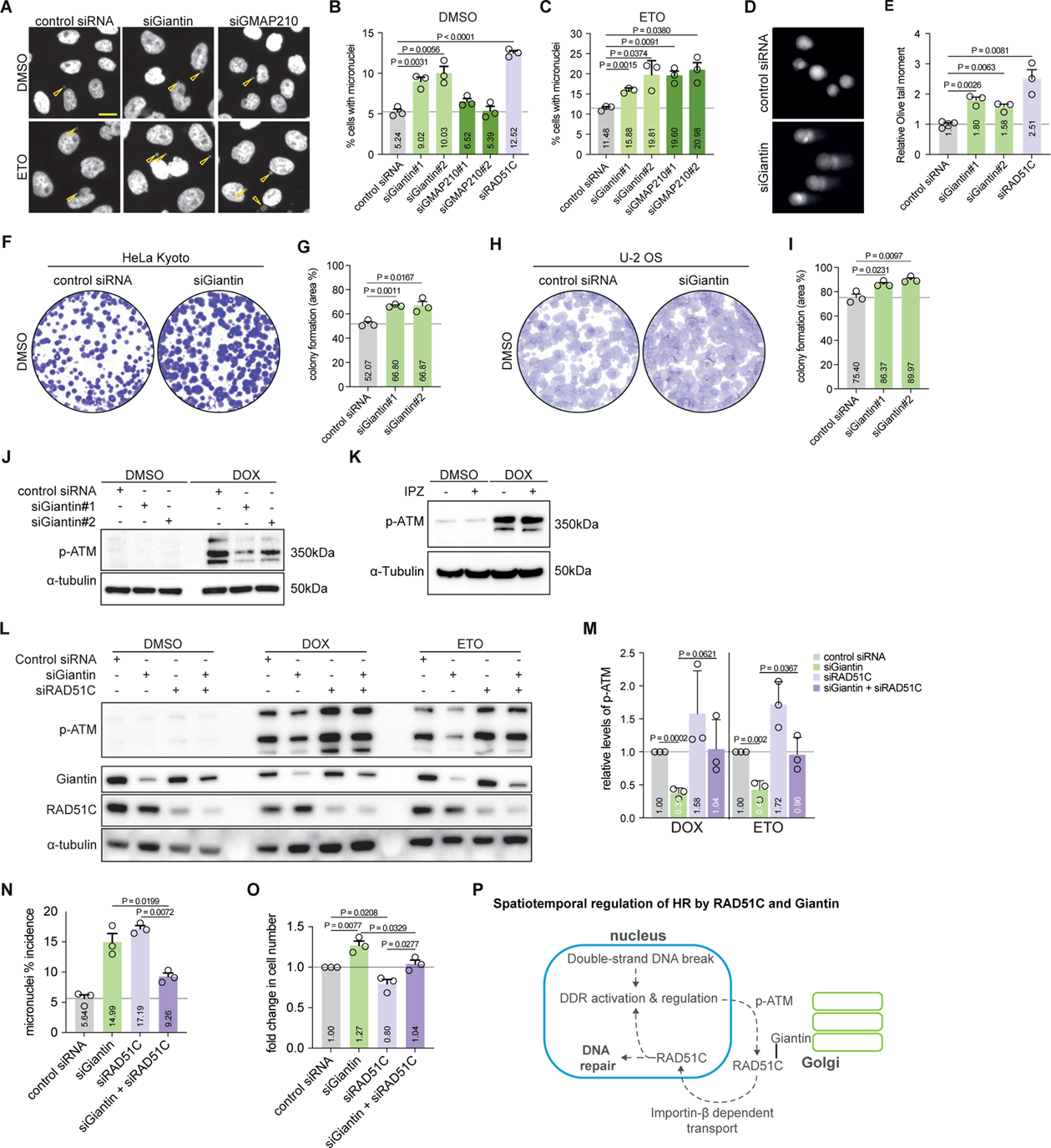
Spatial dysregulation of RAD51C by depletion of Giantin leads to genomic instability, inhibition of DDR signalling and accelerated cell proliferation. (**A–C**) Representative images and associated quantification of micronuclei formation following siRNA treatment under (**B**) basal condition and (**C**) after etoposide-induction DNA damage (10 μM; 16 h followed by 6 h recovery); yellow arrows denote micronuclei structures. Data represent the mean ± s.e.m. (n = 3 biologically independent samples with >10,000 cell analysed). (**D**) Representative comet assay images measuring genomic DNA fragmentation in HeLa-K cells treated with control, Giantin, or RAD51C targeting siRNA. (**E**) Quantification of OliveMoment from comet assay. Data represent the mean ± s.e.m. (n = 3 biologically independent samples). (**F–I**) Representative colony formation assays in (**F**) HeLa-K and (**H**) U-2 OS cell lines with corresponding quantification of colony area coverage in (**G**) HeLa-K and (**H**) U-2 OS cells. Statistical significance was determined using a two-tailed unpaired Student’s t-test. Scale bars, 10 μm. (**J**) WB analysis of HeLa-K cells transfected with control and Giantin siRNA and treated with DOX (40 μM; 3 h). Extracts were prepared and immunoblotted as indicated (n = 2 biologically independent samples). (**K**) WB of HeLa-K cells treated with DOX (40 μM; 3 h) and IPZ (20 μM; 3 h). (**L**) WB of HeLa-K cells transfected with siRNAs against Giantin, RAD51C or a combination of both followed by treatment with DMSO, DOX (40 μM; 3h) or ETO (10 μM; 3 h) (**M**) with associated quantification showing the relative levels of pATM; (n=3 biological independent experiments), data represent the mean ± s.e.m. (**N and O**) Quantification of micronuclei formation and fold change in cell number after siRNA treatments (mean ± s.e.m., n = 3, ≥1,000 cells per condition). (**P**) Proposed model for the regulation of HR-mediated repair through the activation of RAD51C at the Golgi complex. RAD51C, a regulatory HR protein, is anchored to the Golgi through its interaction with the cytoplasmic tail of Giantin, in response to double-strand DNA breaks, this RAD51C Golgi population redistributes to form nuclear foci. This response requires Importin-beta-mediated nuclear import and the phosphorylation of ATM protein kinase. We propose that the Golgi functions as a spatiotemporal timing module that gates RAD51C (and other DDR factors) availability to ensure proper homologous recombination and DNA-damage signaling.

To corroborate basal genomic instability, we performed comet assays (Møller 2018) to measure levels of fragmented genomic DNA (**Figures 5D and 5E**). Cells with defective DNA repair or exposed to DNA-damaging agents display long comet tails, whereas healthy cells show shorter or no tails. Knockdown of Giantin or RAD51C resulted in notably longer comet tails compared to control siRNA treatment (**Figure 5D**). Quantification revealed a 2.5-fold increase in tail length and fluorescence intensity for RAD51C-depleted cells, and 1.8 to 1.6- fold increases for Giantin-depleted cells (siGiantin#1 and siGiantin#2, respectively) (**Figure 5E**).

We next examined the long-term consequences on cell proliferation and colony formation in HeLa-K and U-2 OS cells (**Figures 5F–5I**). Giantin knockdown significantly enhanced both proliferation and colony formation. To determine whether these effects were associated with cell cycle changes, we analysed the cell cycle profiles of Giantin-depleted cells via FACS (**Figure S9D**). No significant cell cycle alterations were detected, suggesting that the increased proliferation is not due to major cell cycle perturbations.

Having observed that Giantin knockdown increases basal genomic instability and displaces RAD51C from the Golgi, we next assessed ATM activation as a proximal readout of HR-linked DDR signalling. To explore this, we assessed the ability of cells depleted of Giantin to initiate HR by measuring ATM phosphorylation, the master regulator of this pathway (**Figures 5J**). In control siRNA treated cells, DOX treatment increased phosphorylated ATM levels in response to DSBs. However, Giantin-depleted cells (**Figure 5J**) showed significantly lower ATM phosphorylation levels under the same conditions.To investigate whether the redistribution of HR factors from the Golgi to the nucleus plays a role in ATM signalling, we used Importazole (IPZ) to inhibit Importin-β-mediated nuclear import and subsequently induced DSBs with DOX (**Figure 5K**). Here, we found that IPZ treatment did not significantly alter ATM signalling, regardless of the induction of DSBs. These findings suggest that the regulated redistribution of HR factors is less impactful on ATM signalling than their mislocalisation, implying that Giantin’s spatial regulation of HR factors is crucial for proper ATM activation.

We next asked whether perturbing RAD51C could modulate the signalling and stability phenotypes observed with Giantin loss. To this end, we carried out co-depletion of RAD51C and Giantin, along with single depletions (**Figures 5L and 5M**). Consistent with previous observations, Giantin knockdown led to a significant decrease in ATM phosphorylation levels, while RAD51C knockdown increased ATM phosphorylation. Co-depletion of RAD51C and Giantin restored ATM phosphorylation levels to those comparable to control cells. To validate the restoration of DDR signalling through co-depletion, we assessed genomic stability via micronuclei incidence (**Figure 5N**). Individual knockdowns of Giantin and RAD51C elevated micronuclei formation, indicative of genomic instability, whereas co-depletion significantly reduced micronuclei incidence, approaching levels observed in control cells. Lastly, we evaluated cell proliferation (**Figure 5O**). While Giantin knockdown enhanced cell proliferation and RAD51C knockdown diminished it, co-depletion resulted in proliferation rates comparable to control. These results are consistent with RAD51C contributing to the Giantin dependent timing mechanism, while not excluding roles for additional Golgi associated factors.

## Discussion

In this study, we reveal evidence for a previously underappreciated and functionally meaningful spatiotemporal regulatory system connecting the Golgi complex and the nucleus, with broad potential implications for homologous recombination (HR) and other DNA repair pathways. Our analysis of Human Protein Atlas (HPA) localisation data (Thul et al. 2017) and experimental validation via siRNA knockdowns uncovered numerous double-localising Golgi-nuclear proteins. Among these, we identified a cluster of DNA repair proteins that exhibit steady-state localisation in both compartments. This cluster encompasses crucial regulatory proteins involved in various DNA repair pathways (**Figure 1D**), not only specific for HR-mediated DNA repair but also Mismatch Repair (MMR), Base Excision DNA Repair (BER) and Microhomology-Mediated End Joining (MMEJ), as well as other integral regulators of DNA repair cellular responses such as chromatin cohesion, ubiquitination, cell cycle regulation and signalling. Taken together, these observations support the existence of a DNA repair protein network that localises to both the Golgi complex and the nucleus, positioning the Golgi as a coordination node that links cytoplasmic organisation with nuclear repair programmes.

Our systematic analysis revealed enrichment of subsets of these DNA repair proteins in distinct sub-Golgi regions. This suggests a compartmentalised platform where components of related DNA repair pathways gather, which could facilitate protein–protein interactions and tune the timing and specificity of downstream signaling. The enriched association of the Golgi scaffold Giantin with HR factors suggests a structural or scaffolding role in organising these repair proteins. These Golgi-resident proteins may stabilise DDR factors at the Golgi and help regulate their release to the nucleus. The close association between Golgins and DNA repair kinases, such as ATM, supports the notion that the Golgi may provide a spatial context for phosphorylation and assembly events. Such spatial organisation could help preserve pathway fidelity by discouraging premature activation and by supporting timely recruitment of DNA damage response components.

In response to genotoxic stress, we observed dynamic shuttling of DNA repair factors between the Golgi and the nucleus, dependent on the type of DNA damage. The correlation between steady-state Golgi localisation and their redistribution patterns upon DNA damage is consistent with a role for the Golgi in temporally coordinating DDR. We propose that the Golgi clusters DDR proteins during steady-state conditions and releases them to the nucleus when damage is detected. In this model, spatiotemporal partitioning functions as a potential checkpoint that coordinates repair factor dosage and timing, thereby integrating cytoplasmic signals with nuclear lesions. Analogous behaviour has been described for shuttling proteins such as BRCA1, whose movements between nucleus and cytoplasm modulate DNA repair, cell cycle regulation and apoptosis (Rodriguez et al. 2004; Feng et al. 2004; Thompson 2010). By extension, Golgi mediated spatiotemporal control of multiple factors may contribute to maintaining genomic stability.

To explore this spatiotemporal regulation, we outline a pathway linking the Golgi to HR repair **(Figure 5P)**. Based on our findings and consistent with established models, we propose that the pathway begins with the detection of double-strand DNA breaks (DSBs) in the nucleus by the MRN complex, which activates ATM kinase. Although canonical MRN function is indispensable for initiating ATM activation at DNA breaks, our data indicate that once ATM is activated in the nucleus, an ATM-dependent signal or possibly a fraction of active ATM can subsequently access the Golgi (Ovejero et al. 2023). In this view, the Golgi acts downstream of the break sensor as an amplification and timing module rather than a primary sensor, representing an organelle level checkpoint that conditions release on damage context, akin to cytoplasmic ATM pools observed in oxidative stress responses (Shiloh and Ziv 2013). Consistent with this model, Golgi-associated scaffolding and kinase activities would be expected to gate the timed availability of a subset of HR regulators for lesion engagement.

Within this framework, our data places Giantin as a spatial modulator of this Golgi-nuclear timing system. Giantin depletion increases basal genomic instability, accelerates cell proliferation, reduces ATM activation after DSB induction while displacing a RAD51C pool to the nucleus. RAD51C depletion alone elevates genome instability but increases phosphorylated ATM after damage, while co-depletion of RAD51C with Giantin partially restores phosphorylated ATM and genome stability toward control and attenuates the Giantin dependent proliferation increase. These findings are consistent with Giantin acting through RAD51C as one conduit, while likely engaging other additional Golgi localising DNA damage response factors.

RAD51C’s known function can plausibly account for part of the Giantin loss phenotype. As a member of the RAD51 paralog complexes BCDX2 and CX3, RAD51C executes essential roles in stabilising replication forks, promoting RAD51 filament assembly, and resolving Holliday junctions (Rein, Bernstein, and Baldock 2021; Sullivan and Bernstein 2018). In principle, mislocalisation could undermine these processes, for example by altering the recruitment of repair factors and DDR signalling with downstream effects on other factors such as CHK2 (Badie et al. 2009; Somyajit, Subramanya, and Nagaraju 2012). As a central participant in HR, RAD51C orchestrates RAD51 filament formation and subsequent strand invasion steps required for DSB repair (Somyajit et al. 2015; Prakash et al. 2022). If prematurely released to the nucleus or improperly retained at the Golgi, RAD51 filaments may be disorganised or suboptimally assembled, leading to error-prone repair events. RAD51C also influences checkpoint signaling by stabilising replication forks and interacting with ATR–RPA, enabling proper CHK2 phosphorylation and cell-cycle arrest upon DNA damage (Badie et al. 2009; Berti et al. 2020). Dysregulation of these checkpoint pathways could encourage cells to progress through the cell cycle despite harboring DNA lesions and alter DDR signalling.

Beyond RAD51C, our screen identified other DDR regulators with dual Golgi–nuclear localisation, notably the HR regulator NBN, and candidates spanning chromatin regulation, ubiquitination and checkpoint signalling. These observations argue for a cohort model in which multiple Golgi-proximal factors are clustered at steady state and released with tuned timing, such that their combined availability, rather than any single protein alone shapes HR pathway behaviour and replication-stress responses. In this view, the Golgi provides a spatiotemporal checkpoint that modulates factor dosage and order of arrival at lesions. Our dataset supports this timing perspective through changes in upstream signalling and genome-stability metrics, but does not dissect how HR repair efficiency is impacted; any effect on HR therefore remains a testable prediction. Systematic follow-up should broaden the catalogue of Golgi-associated DDR factors, benchmark their relative contributions, and couple temporal proteomics with multiplex perturbations to resolve how joint release kinetics influence HR fidelity and fork protection and restart.

In a clinical context, perturbation of Golgins, including Giantin, has been linked to elevated breast cancer risk (Wang et al. 2020; Craven, Gökmen-Polar, and Badve 2021; Mathioudaki et al. 2021; Pipek et al. 2023; Ghannoum et al. 2023; Leighton et al. 2023), as well as decreased patient survival (**Figures S9E and S9F**) (Győrffy 2021; Ősz, Lánczky, and Győrffy 2021). Together with reports implicating additional Golgin family members across various cancers (Baschieri et al. 2015; Bhat et al. 2017), these associates are compatible with a broader role for Golgi scaffolding and signaling processes in controlling genomic integrity. This raises the prospect of therapeutically modulating organelle-level checkpoints either by tuning Golgi-dependent timing of factor release or by leveraging Golgi–nuclear communication to sensitise tumours to genotoxic agents.

Finally, the observation that many other proteins dually localise to the Golgi and nucleus, spanning membrane trafficking, RNA metabolism, and lipid regulation, supports the concept of the Golgi as a dynamic integration hub for multiple essential pathways. Its proposed role in DNA repair orchestration may extend beyond HR to additional DDR networks. Altogether, these findings motivate a shift from purely concentration-based models of the DDR to timing-aware, compartment-resolved models, opening new avenues for both fundamental research and translational approaches in DNA repair and cancer biology.

## Materials and Methods

### Antibodies and chemicals

Several commercially antibodies and chemical were used in the paper, including RAD51C (ab72063, Abcam; 1:500 for immunostaining) & (ab95069, Abcam, 1:2000 for western blotting; 1:500 for immunoprecipitation), GOLGB1/Giantin (AF8159, R&D systems; 1:500 for immunostaining and immunoprecipitation, 1:2000 for western blotting) & (G1/M1, Enzo Life Sciences, 1:500 for immunostaining), GM130 (610822, BD Biosciences; 1:500 for immunostaining, 1:2000 for western blotting), TGN46 (AHP500GT, Bio-Rad; 1:1000 for immunostaining; 1:2000 for western blotting), ATM (MA1-23152, Invitrogen; 1:2000 for western blotting), pATM Ser1981 (MA1-2020, Invitrogen; 1:500 for immunostaining; 1:2000 for western blotting), CHK2 pThr68 (PA5-17818, Invitrogen; 1:2000 for western blotting), DNA-PKcs (Ser2056) (PA5-78130, Invitrogen, 1:2000 for western blotting), gamma-H2AX pSer139 (613402, Biolegend; 1:500 for immunostaining, 1:2000 for western blotting), alpha-tubulin (MS-581, Thermo Fisher; 1:10,000 for western blotting) Lamin B1 (ab16048, Abcam; 1:2000 for western blotting), doxorubicin (ab120629, Abcam); Importazole (SML0341, Sigma-Aldrich), KU55933 (ATMi, SML1109, Sigma-Aldrich); ve-821 (ATRi, HY-14731; MedChemExpress) NU7441 (DNA-PKi, HY-11006; MedChemExpress), etoposide (ab120227, Abcam), mitomycin C (M7949, Sigma-Aldrich), hydrogen peroxide (H1009, Sigma-Aldrich), potassium bromate (104912, Merck Millipore), camptothecin (ab120115, Abcam), SYBR Gold (S11494, Invitrogen), crystal violet (C0775-100G, Sigma-Aldrich).

### Cell lines, cell culture and siRNA transfection

HeLa Kyoto, U-2 OS, MCF7, were cultured in DMEM (Life Technologies) supplemented with 10% FBS (Invitrogen) and 1% L-glutamine (Invitrogen). Cells were checked for mycoplasma contamination by PCR. siRNA transfections were performed with Lipofectamine 2000 (Invitrogen) using SilencerSelect siRNAs (Ambion) according to the manufacturer’s instructions. Transfections were carried out for 72 h and the final siRNA concentrations used were 15nM for all siRNAs. Giantin siRNA-1: 5951 & siRNA-2: 5953; GMAP210 siRNA-1: s17811 & siRNA-2: s17812; RAD51C siRNA: s11737.

### Localisation validation siRNA assay

A custom-designed siRNA library targeting our proteins of interest (Ambion) was designed and prepared in 96-well glass bottom plates (Miltenyi Biotec) using a protocol for solid phase reverse transfection as previously described (Erfle et al. 2008; Stadler et al. 2012). Non-targeting siRNA was used as a negative control. 72 hours after cell seeding, cells were fixed with 4% paraformaldehyde (PFA), permeabilised with 0.1% Triton-X100 and immunostained against the HPA antibodies of interest and a Golgi marker, GM130. Hoechst 33342 was used as a nuclear stain. siRNAs and antibodies details are available in Supplementary Table S1. Images were acquired on a fully automated Molecular Devices IXM with a 10x/0.45 NA P-APO objective. The resulting images were analysed using Cell Profiler software (Carpenter et al. 2006) for quantitative and automated measurements of fluorescence from the antibodies as previously described (Stadler et al. 2012). Briefly, nuclei were segmented in the nuclear channel, the Golgi complex was segmented in the Golgi marker channel and using the segmented nuclei as seeds, the two structures were associated. Intensity profiles of each compartment were acquired. A reduction of 25% or more of the antibody staining in both Golgi and nuclear compartments was considered as a validation of the antibody specificity **(Figures S1 and S2; Supplementary Table S1)**.

### Golgi cisternal localisation analysis

As previously described (Dejgaard et al. 2007), HeLa-K cells were treated with nocodazole (33 μM) or the solvent control methanol for 3 h. After treatment, cells were fixed with 4% PFA, permeabilized with 0.1% Triton-X100 and immunostained using antibodies against *trans*-Golgi marker, TGN46, *cis*-Golgi marker, GM130 and the protein of interest. Images of the Golgi mini-stacks were acquired using a confocal microscope Olympus Fluoview FV3000 with a 60x/1.3 NA silicon oil immersion apochromatic objective. Golgi mini-stacks were visually inspected, and those that were in the same plane were selected for analysis. Images were analysed using Fiji (Schindelin et al. 2012); the relative position of the fluorescence profile of the protein of interest against the *trans*- and *cis-Golgi* markers was measured using a plot profile tool. The measured distances were used to calculate Pearson’s correlation coefficient (PCC) between *cis*-Golgi and *trans*-Golgi markers and proteins of interest.

### Drug treatments

Cells were treated 24 h after seeding. For camptothecin, etoposide and mitomycin C experiments, cells were treated for 16 h at a concentration of 0.1 μM, 50 μM and 5 μM, respectively. After treatment cells were incubated with fresh medium for 2 hours. H_2_O_2_ experiments were performed by treating cells for 20 min at a concentration of 50 μM, followed by change with fresh medium and 15 min recovery. For nocodazole and KBrO_3_ experiments cells were treated for 3 h at the concentration of 33 μM and 5 mM, respectively. Doxorubicin was used at a concentration of 40 μM for 3 h unless indicated otherwise. For kinase and importin-β inhibitor treatments, cells were pretreated for 30 min with the described inhibitor prior to the addition of doxorubicin. The inhibitors were used at the following concentrations: importazole (IPZ) (20 μM), KU55933 (ATMi) (30 μM), NU7441 (DNA-Pki) (10 μM) and ve-821 (ATRi) (10 μM).

### Immunofluorescence assay

Cells were fixed with 4% paraformaldehyde in PBS and permeabilised with 0.1% Triton X-100 for 15 min at room temperature, then cells were blocked with 5% bovine serum albumin in 0.05% Triton X-100 for 60 min and incubated with primary antibodies in blocking buffer at room temperature for 3 h. Following 3 washes with PBS, cells were incubated with fluorescent dye-conjugated secondary antibodies diluted in a blocking buffer for 1 h at room temperature.

### Image and data analysis

Confocal microscopy experiments were performed with fixed and immunostained cell samples on a microscope FV3000, Olympus. Z stacks of images covering the entire cell thickness were acquired. All image analysis was performed using Cell Profiler (Carpenter et al. 2006) and Fiji (Schindelin et al. 2012). Briefly, first nuclei were segmented in the Hoechst channel. When appropriate, the Golgi complex was segmented in the Golgi marker channel and using the segmented nuclei as seeds, the two structures were associated. Intensity profiles, morphology features and structuring counting analysis were performed when required using Cell Profiler.

### Comet assay

The assay was carried out as previously described (Vodenkova et al. 2020). Briefly, cells were trypsinised, pelletised and resuspended in ice-cold PBS at a concentration of 25 × 10^4^ cells per ml. The cells were resuspended in 2% low melting agarose (Sigma) and spread quickly onto gel bond film (Biozym) covered in 1% agarose (Sigma). Samples were immersed into a lysis buffer (100 mM EDTA, 2.5 M NaCl, 10 mM Tris-HCl and 1% Triton-X100; pH 10) overnight at 4°C. Followed by a wash with ice-cold water and run in an electrophoresis chamber (alkaline buffer: 1 mM EDTA, 300 mM NaOH; pH 13) at 15 V, 300 mA for 60 min at 4°C. Slides were first washed in a Tris-HCl neutralisation buffer (0.4 M; pH 7.5) followed by water, stained with SYBR Gold (Thermo Fisher Scientific) (1:10000) and finally dried. Comets were imaged by an automated Olympus Scan^R screening microscope and comet tails were scored using OpenComet plugin (Gyori et al. 2014).

### Western blotting analysis

HeLa-K cells were lysed using RIPA buffer (Thermo Scientific) with a complete protease inhibitor cocktail (Roche). SDS-PAGE was performed on pre-cast Tris-Acetate gels followed by transfer to PVDF transfer membrane (Merck Millipore). Proteins were detected using primary antibodies as described followed by incubation with secondary antibodies coupled with HRP (Invitrogen). Detection of protein was performed using Pierce ECL Plus Western Blotting Substrate reagent (Thermo Scientific) and visualised on Azure 280 chemiluminescent imaging system. Golgi enrichment assay was performed using Minute™ Golgi Apparatus Enrichment Kit (GO-037, Invent Biotechnologies) according to the manufacturer’s instructions. The purity of the cis and trans-Golgi fractions was confirmed by using Golgi markers GM130 for the *cis*-Golgi and TGN-46 for the *trans-*Golgi. Subcellular fractionation was performed using a subcellular protein fractionation kit for cultured cells (78840, Thermo Scientific) according to the manufacturer’s instructions. Fractions were verified using well-established markers (GM130 for Golgi membranes, Lamin B1 for the nuclear compartment, and alpha-tubulin for the cytoplasmic fraction) and probed for proteins of interest presence in each compartment. Protein membrane:nuclear distribution ratio was calculated by first dividing the protein level in each fraction by the protein level of the appropriate control, the resulting membrane (M_protein_ / M_GM130_) and nuclear (N_protein_ / N_laminB1_) ratios were further divided to give the ratio = (M_protein_ / M_GM130_) / (N_protein_ / N_laminB1_). The ratios obtained from the control were normalised to 1 and compared to the treated group.

### Immunoprecipitation

HeLa-K cells were lysed using a lysis buffer (50 mM Hepes, 130 mM NaCl, 1 mM DTT 1% NP-40) with a complete protease inhibitor cocktail (Roche). Cell lysates were centrifuged at 13,000 rpm for 10 mins at 4°C. For immunoprecipitation, the lysates were incubated with the primary antibody described and rotated overnight at 4°C. Next, the lysates were incubated with G-agarose beads (Roche) and rotated for 4 h at 4°C. The samples were washed with cold lysis buffer and then precipitated proteins were eluted by 2x SDS sample buffer and analysed by Western blotting.

### Colony formation assay

HeLa-K or U-2 OS cells, transfected with either a Giantin or control siRNA were seeded on a 6-well plate (500 cells/well) and allowed to grow for 12 days in a complete culture medium. Where indicated, 24h after seeding the cells were treated with a DNA damaging agent, DOX (1 μM) or a solvent control, DMSO for 24h. Subsequently, the medium was replaced, and it was refreshed every 4 days. The resulting colonies were washed twice with PBS, followed by fixing and staining for 30 min with a solution containing 0.1% (w/v) crystal violet and 20% (v/v) ethanol. Then the colonies were washed again to remove excess crystal violet with PBS. Finally, after the dishes were dry, digital images of the colonies were acquired using a camera and quantified.

### Statistical analysis

All data were obtained from at least three independent experiments if not otherwise stated. Statistical analyses were performed using a two-tailed t-test for pairwise comparison and one-way analysis of variance (ANOVA) for multiple comparisons on GraphPad Prism 9. Data are expressed as the standard error of the mean (s.e.m.). n values indicate biologically independent samples and experiments. P < 0.05 was considered statistically significant.

## Acknowledgements

We thank the ALMF, FCCF and the Pepperkok team for their support. In addition, we acknowledge Jan Ellenberg, Claudia Lukas, Alba Diz-Muñoz, Diana Ordonez, Per Haberkant and Simone Köhler for their assistance in developing the project and manuscript. G.G. was supported by the EMBL EIPOD programme, K.K. by the EMBL PhD programme and M.M.K. by the German Centre for Lung Research.

---

**Figure S1:**
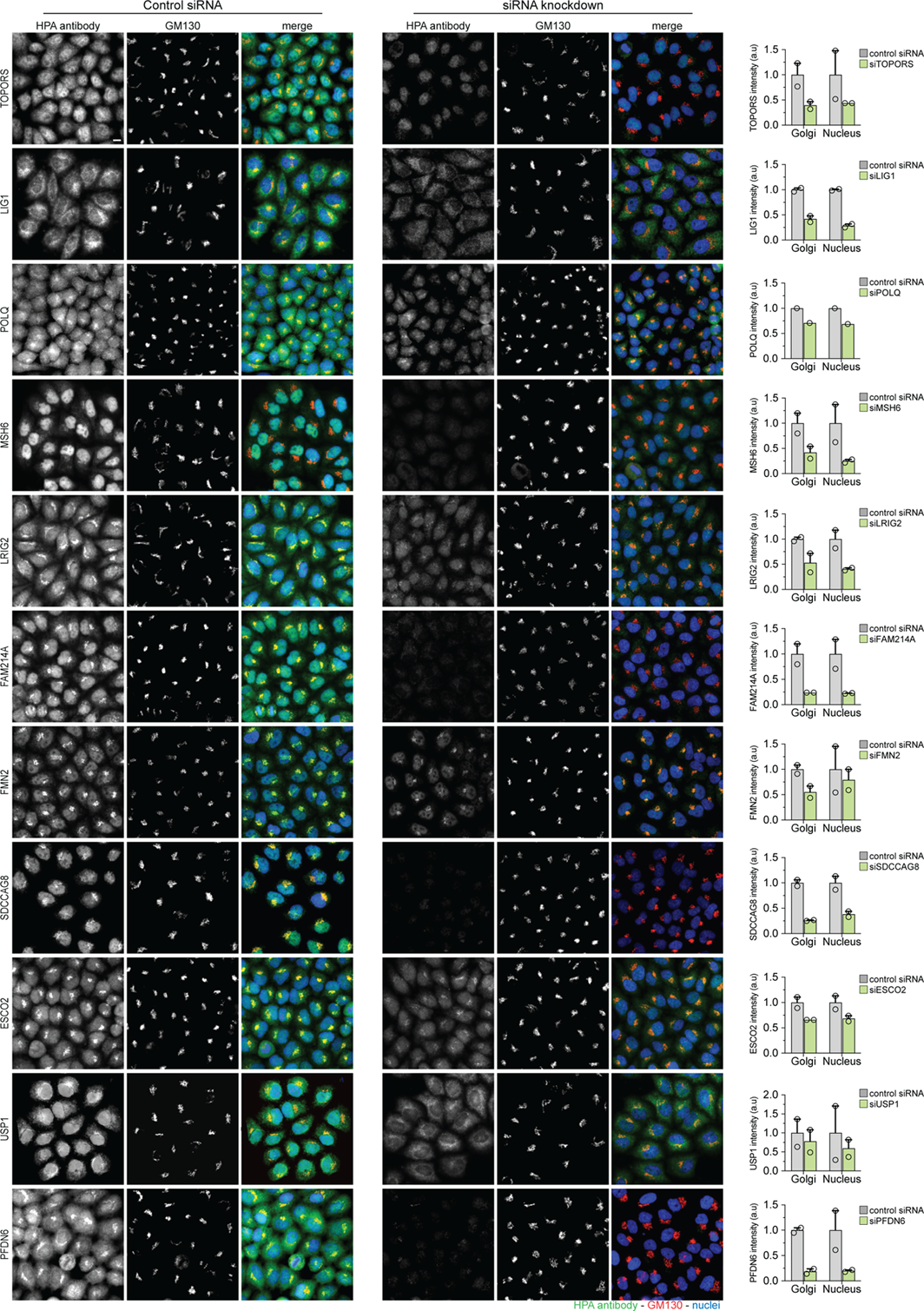
Validation of DDR protein antibody specificity and localisation, related fo Figure 1. Representative images of HeLa-K cells stained with antibodies against DDR proteins and the Golgi marker, GM130. Cells were treated with a control or an siRNA against the protein of interest to validate the specificity of the antibody. The changes in intensity measurements in the Golgi and nuclear compartment were quantified (n=2 for all proteins except for POLQ, which had n=1, biological independent samples with at least 500 cells analysed for each treatment). Scale bar, 10 μm. Continued on **Figure S2A**.

**Figure S2:**
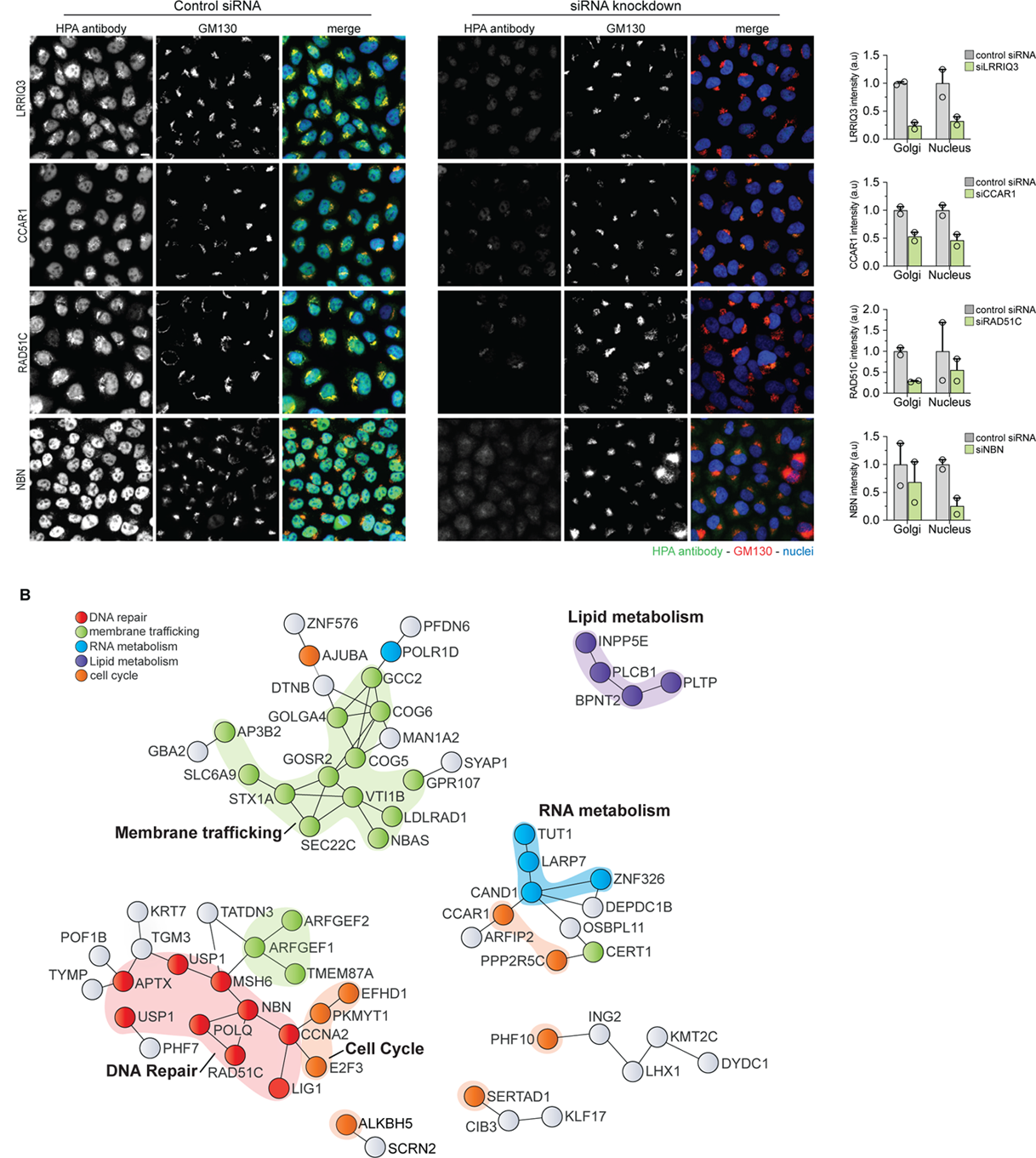
Validation of DDR protein localisation and functional network analysis, related to Figure 1. **(A)** Representative images of HeLa-K cells stained with antibodies against DDR proteins and the Golgi marker, GM130. Cells were treated with a control or an siRNA against the protein of interest to validate the specificity of the antibody. The changes in intensity measurements in the Golgi and nuclear compartment were quantified. (n=2, biological independent samples with at least 500 cells analysed for each treatment). **(B)** Experimentally-based protein-protein interaction networks displaying the Golgi-nuclear localisation proteins validated in this study. Proteins are categorised by functional pathways annotated using gene ontology terms **(Table S1)**.

**Figure S3:**
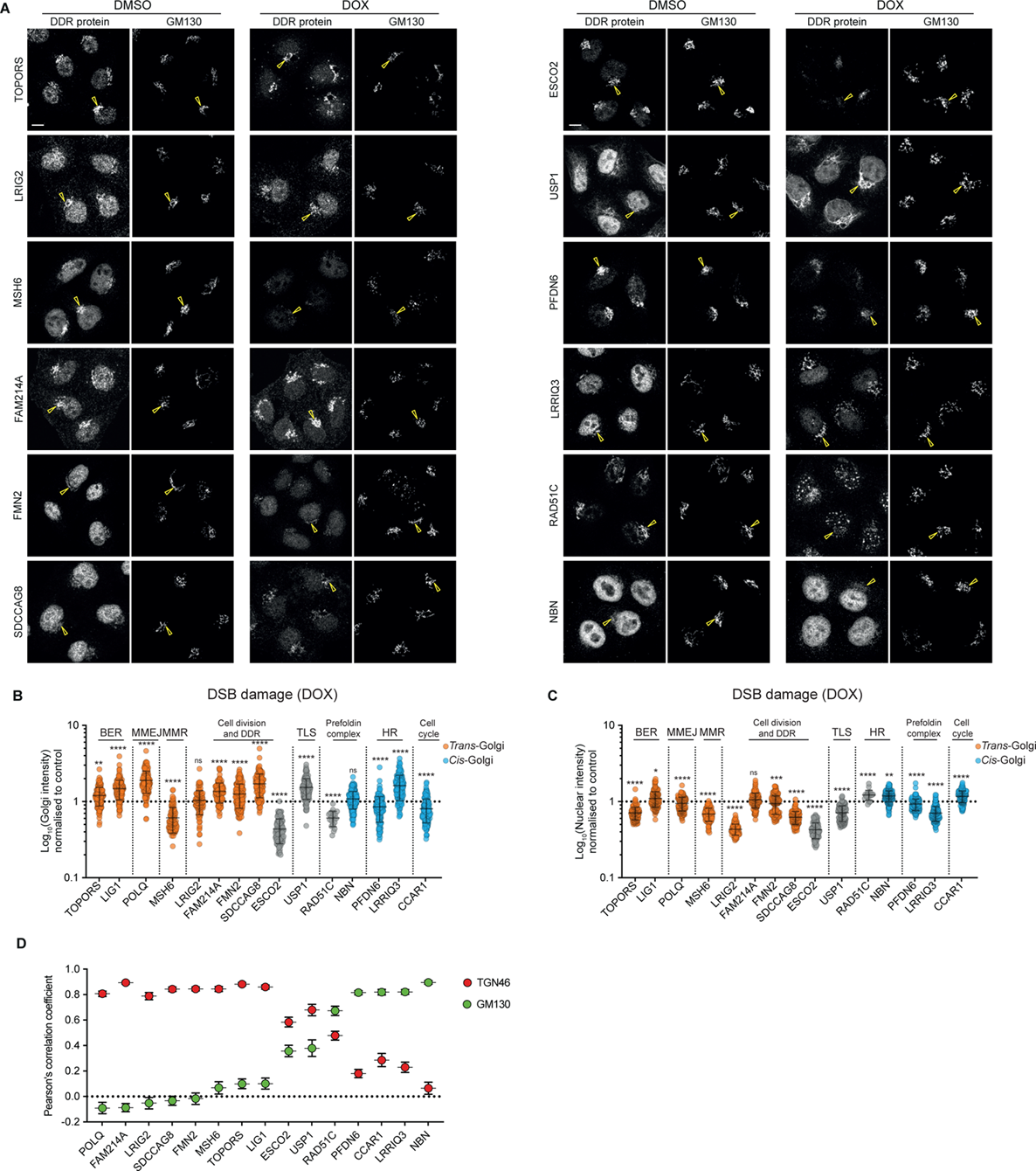
DOX-Induced Redistribution and Golgi-cisternal localisation analysis of DDR proteins, related to Figure 2. **(A)** Representative images of HeLa-K cells stained with antibodies against DDR proteins (green) and the Golgi marker, GM130 (red). Cells were treated with DOX (40 μM, 3 h), fixed, and immunostained. Yellow arrows denote Golgi membranes. Scale bars, 10 μm. **(B and C)** Quantification of DDR protein intensity changes upon DOX treatment. **(B)** Relative intensity of DDR proteins at the Golgi and **(C)** in the nucleus, normalised to untreated control. Data represent the mean ± s.e.m. (n = 3 biologically independent samples with at least 200 cells analysed for each protein in control and treatment conditions). The proteins are classified according to their Golgi localisation patterns (Figure 2E). Statistical significance was determined using a two-tailed unpaired Student’s t-test; ns P > 0.05, *P < 0.05, **P < 0.01, ***P < 0.001, ****P <0.0001, compared to untreated control. **(D)** Golgi-cisternal localisation analysis of Golgi-nuclear DDR proteins. HeLa-K cells were treated with Nocodazole (33 μM, 3 h), fixed and stained with antibodies against DDR proteins. Quantification of Pearson’s correlation coefficient (PCC) between cis-Golgi and trans-Golgi markers and DDR proteins; N ≥ 4 cells; n ≥ 36 mini stacks.

**Figure S4:**
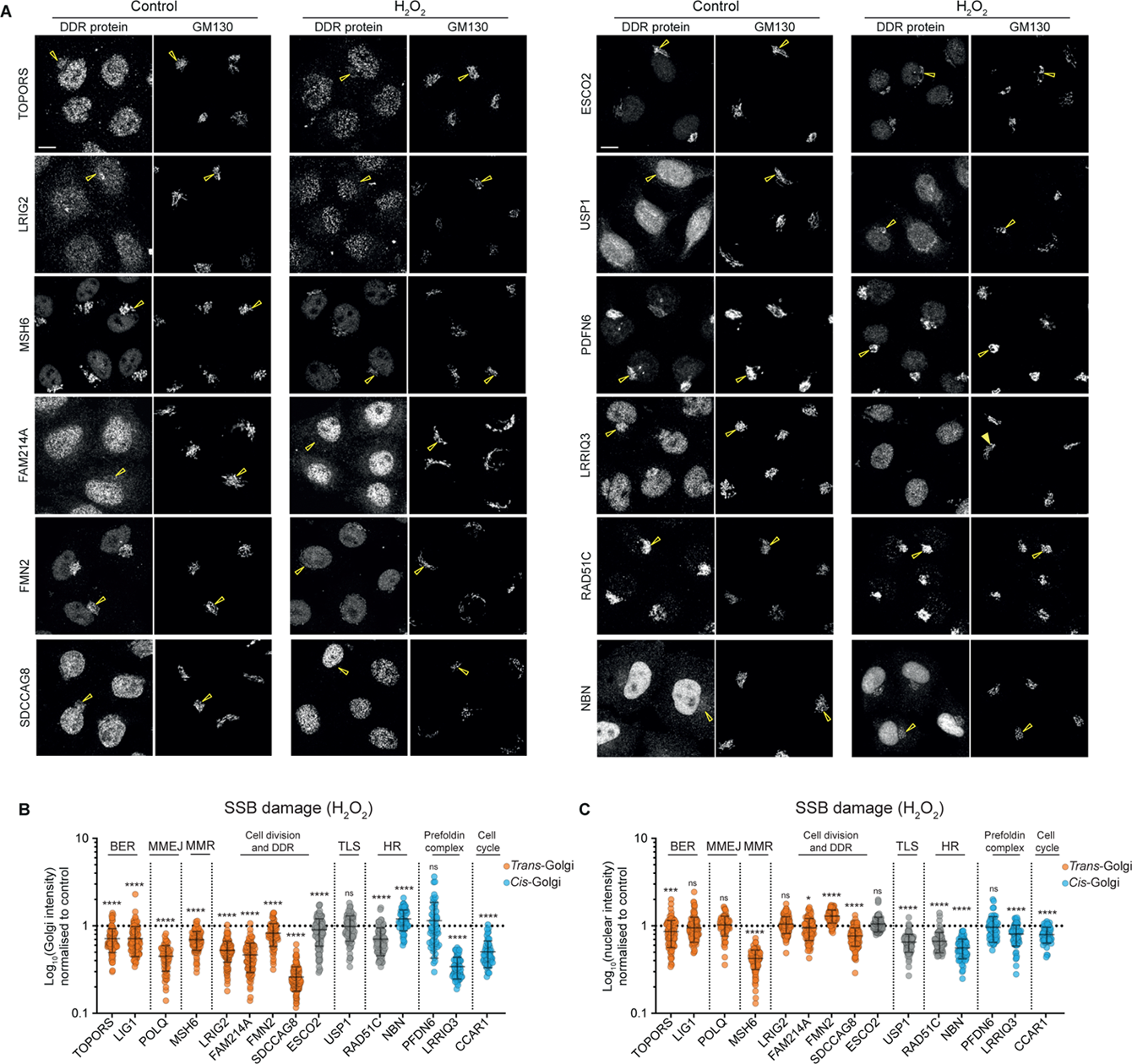
H_2_O_2_-Induced redistribution of DDR proteins, related to Figure 2. **(A)** Representative images of HeLa-K cells were stained with antibodies against DDR proteins and the Golgi marker, GM130. HeLa-K cells were treated with H2O2 (50 μM, for 20 min followed by 15 min recovery), fixed and stained with antibodies against DDR proteins and the Golgi marker, GM130; yellow arrows denote the Golgi membranes. Scale bars, 10 μm. **(B and C)** Quantification of DDR protein intensity changes upon H_2_O_2_ treatment. **(B)** Relative intensity of DDR proteins at the Golgi and **(C)** in the nucleus, normalised to untreated control. Data represent the mean ± s.e.m. (n = 3 biologically independent samples with at least 200 cells analysed for each protein in control and treatment conditions). The proteins are classified according to their Golgi localisation patterns (Figure 2E). Statistical significance was determined using a two-tailed unpaired Student’s t-test; ns P > 0.05, *P < 0.05, ***P < 0.001, ****P <0.0001, compared to untreated control.

**Figure S5:**
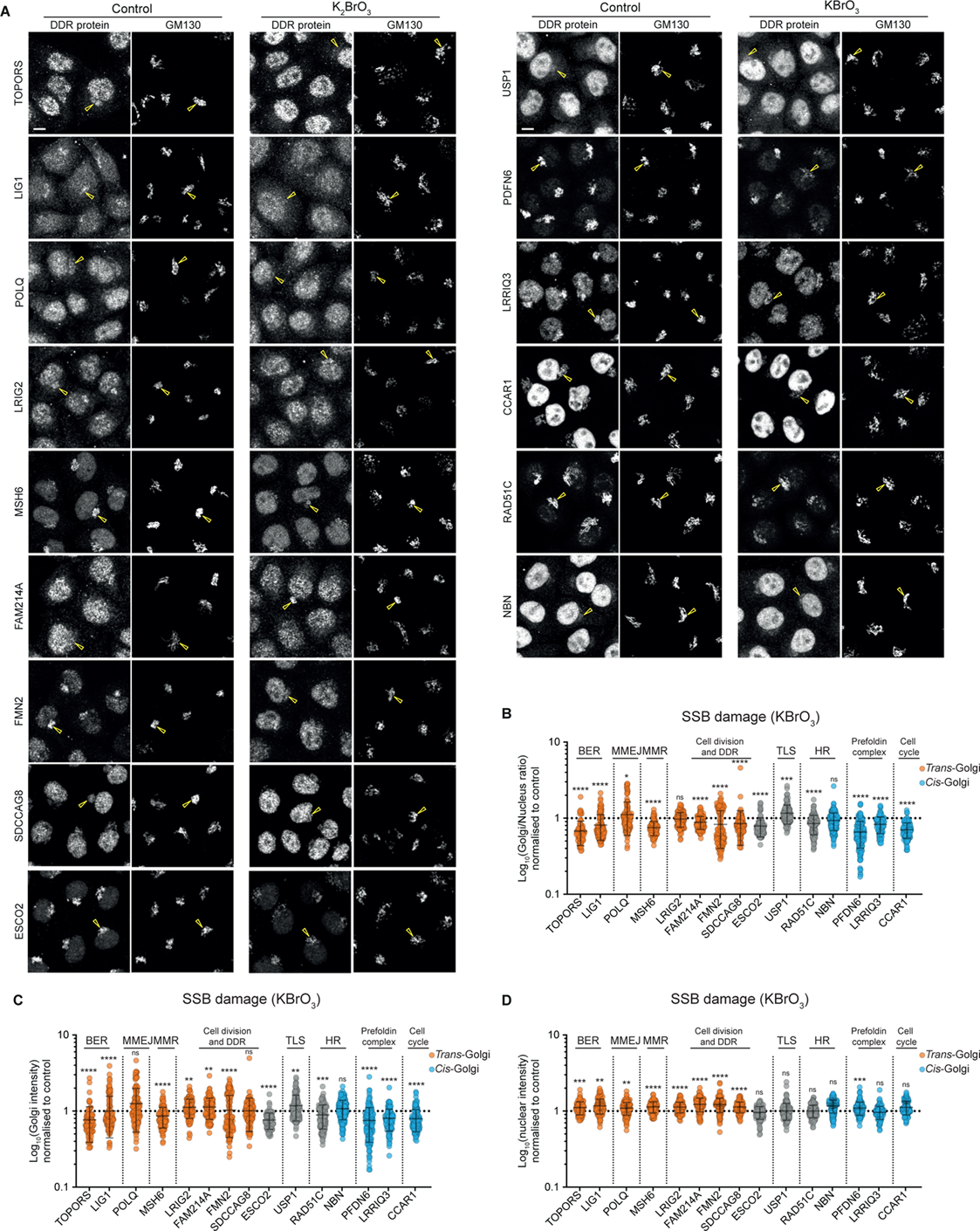
KBrO_3_-Induced redistribution of DDR proteins, related to Figure 2. **(A)** Representative images of HeLa-K cells were stained with antibodies against DDR proteins and the Golgi marker, GM130. HeLa-K cells were treated with KBrO_3_ (5 mM, 3h), fixed and stained with antibodies against DDR proteins and the Golgi marker, GM130 (A to D); yellow arrows denote the Golgi membranes. Scale bar, 10 μm. **(B)** A ratio of DDR protein Golgi-nuclear distribution after treatment with KBrO_3_. **(C and D)** Quantification of DDR protein intensity changes upon KBrO_3_ treatment. **(C)** Relative intensity of DDR proteins at the Golgi and **(D)** in the nucleus, normalised to untreated control. Data represent the mean ± s.e.m. (n = 3 biologically independent samples with at least 200 cells analysed for each protein in control and treatment conditions). The proteins are classified according to their Golgi localisation patterns (Figure 2E). Statistical significance was determined using a two-tailed unpaired Student’s t-test; ns P > 0.05, *P < 0.05, **P < 0.01, ***P < 0.001, ****P <0.0001, compared to untreated control.

**Figure S6:**
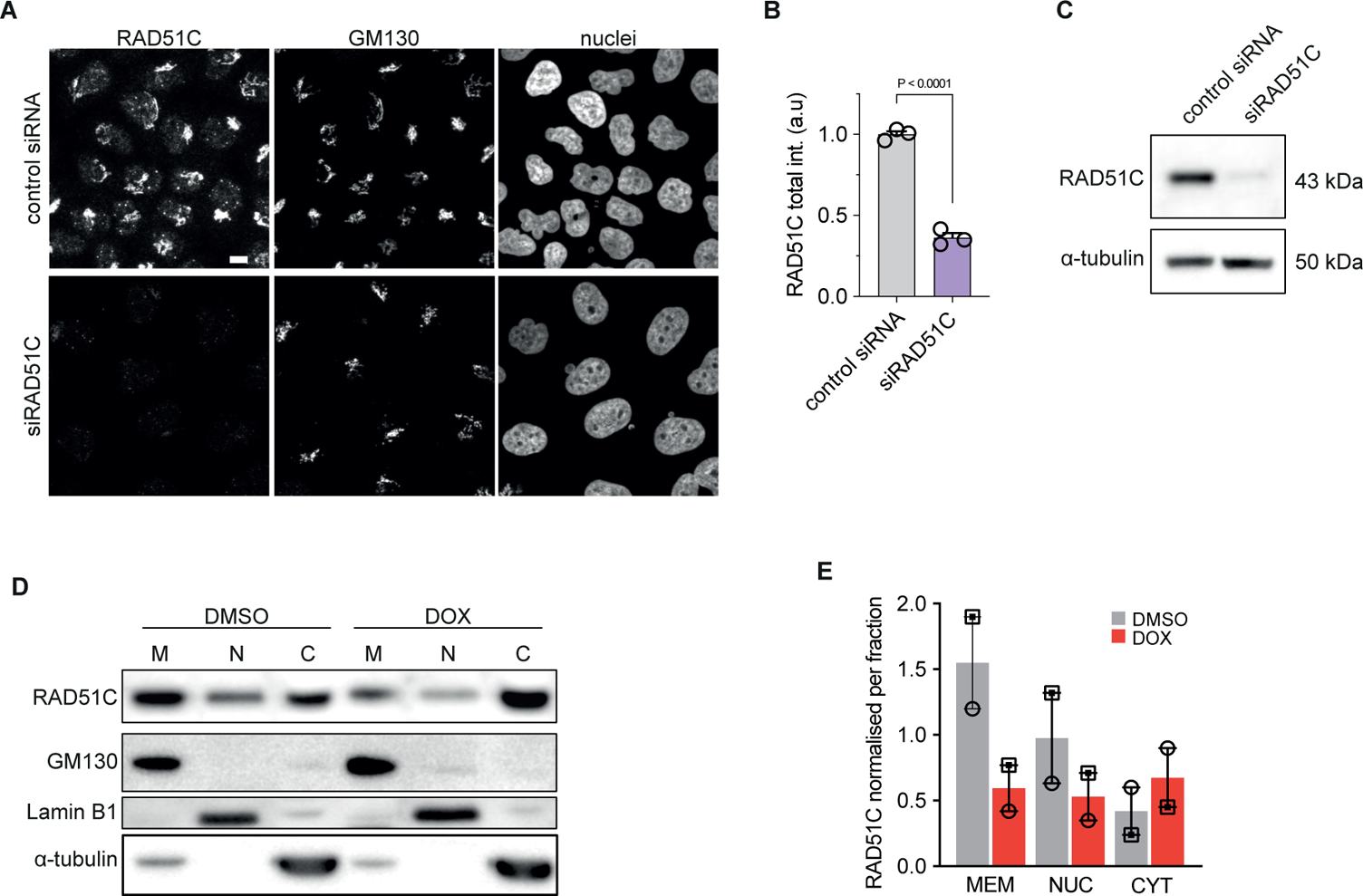
Validation of RAD51C subcellular localisation, related to Figure 3. **(A)** Representative images showing the RAD51C antibody specificity. HeLa-K cells were stained with antibodies against RAD51C and GM130; DNA was stained with Hoechst 33342. Scale bar, 10 μm. **(B)** Quantification of RAD51C sum intensity after RAD51C depletion. Data represent the mean ± s.e.m. (n = 3 biologically independent samples with a total of 831 cells analysed). **(C)** Western blot analysis showing the level of RAD51C protein level, in RAD51C-depleted and control cells (n = 3 biologically independent samples). Statistical significance was determined using a two-tailed unpaired Student’s t-test. **(D)** Western blot analysis showing the subcellular localisation of RAD51C treated with the DMSO and doxorubicin. The protein levels normalised against the respective fraction control (RAD51Cmem/GM130; RAD51Cnuc/Lamin B1; RAD51Ccyt/tubulin). Data represent the mean ± s.e.m; (n = 2 biologically independent samples). **(E)** Quantification of RAD51C membrane-nuclear distribution ratio after treatment with DOX, calculated from isolated fractions.

**Figure S7:**
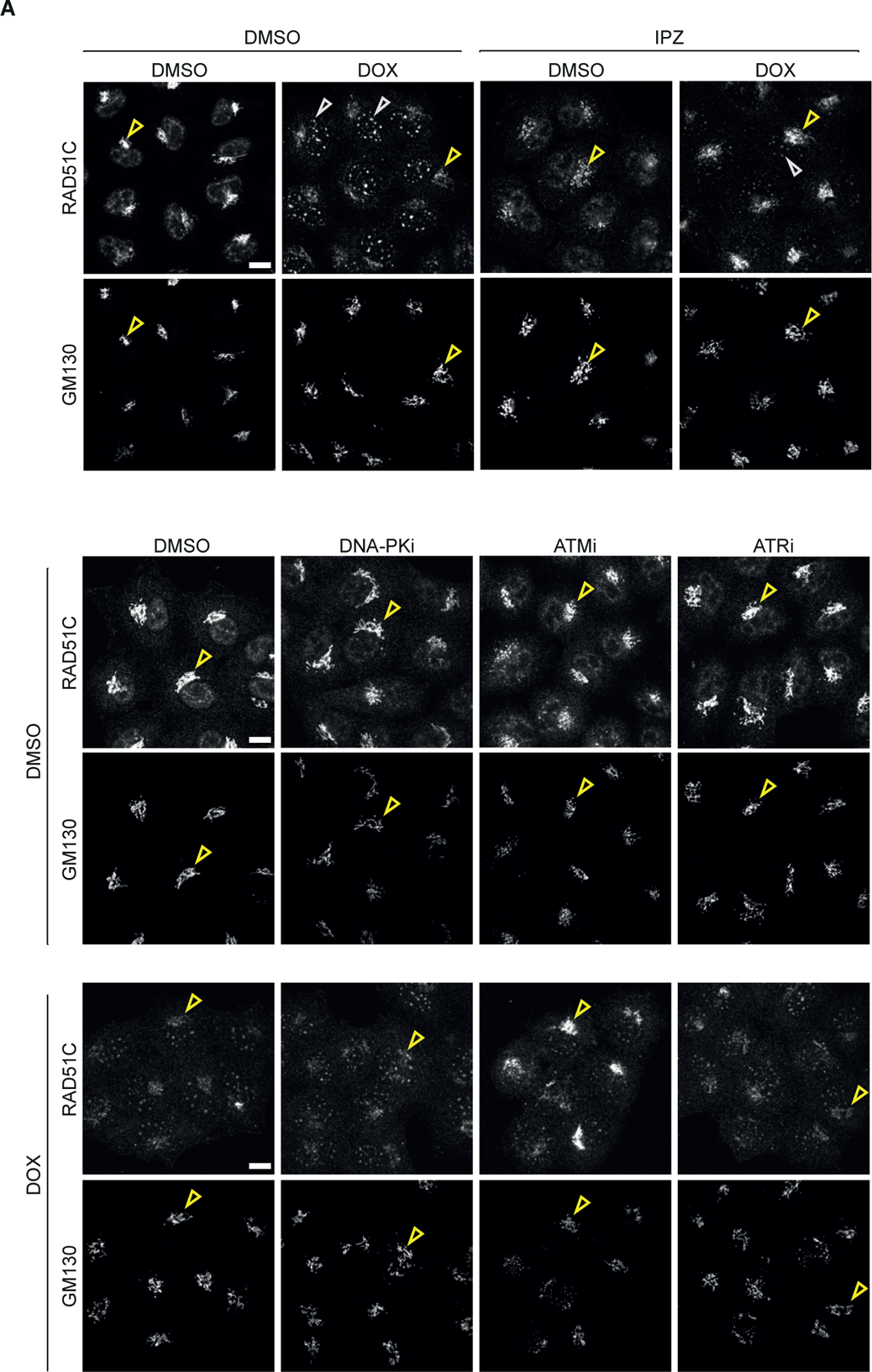
RAD51C Golgi population redistribution upon treatment with IPZ and inhibition of DDR signalling, related to Figure 3. **(A)** Representative images of the RAD51C protein redistribution upon treatment with IPZ and DOX. HeLa-K cells were stained with antibodies against RAD51C and GM130. Cells were treated with DMSO or IPZ prior to a 3-hour treatment with DOX. **(B)** Representative images of the RAD51C protein redistribution upon treatment with DDR signalling inhibitors and DOX. HeLa-K cells were treated with DMSO (control) alone or with ATM inhibitor (KU55933), ATR inhibitor (VE-821) or DNA-PK inhibitor (NU7441) prior to a 3-hour treatment with doxorubicin. Yellow arrows denote the Golgi membrane; white denotes nuclear foci. Scale bars, 10 μm.

**Figure S8:**
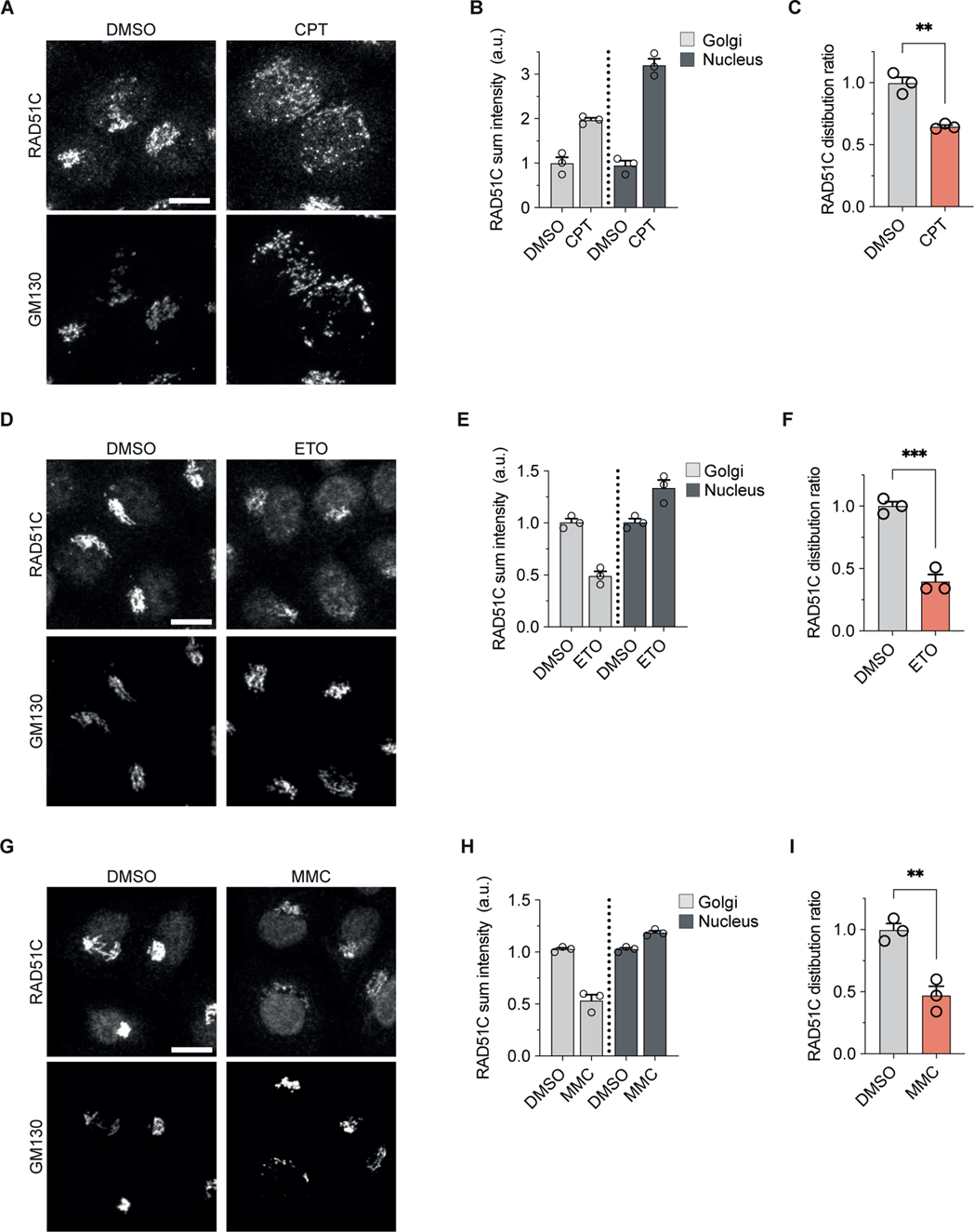
RAD51C protein localisation and redistribution upon induction treatment with DNA damage agents, related to Figure 3. **(A)** Representative images of HeLa-K cells stained with antibodies against RAD51C and GM130 after treatment with Camptothecin (CPT) for 16 h followed by media change for 2 h. Scale bar, 10 μm. **(B)** Quantification of RAD51C distribution between the Golgi and nuclear compartments after CPT treatment. **(C)** Quantification of RAD51C distribution ratio between the Golgi and nuclear compartments after CPT treatment. **(D)** Representative images of HeLa-K cells stained with antibodies against RAD51C and GM130 after treatment with Etoposide (ETO) for 16 hours followed by media change for 2 h. Scale bar, 10 μm. **(E)** Quantification of RAD51C distribution between the Golgi and nuclear compartments after ETO treatment. **(F)** Quantification of RAD51C distribution ratio between the Golgi and nuclear compartments after ETO treatment. **(G)** Representative images of HeLa-K cells stained with antibodies against RAD51C and GM130 after treatment with Mitomycin C (MMC)for 16 h followed by media change for 2 h. Scale bar, 10 μm. **(H)** Quantification of RAD51C distribution between the Golgi and nuclear compartments after MMC treatment. **(I)** Quantification of RAD51C distribution ratio between the Golgi and nuclear compartments after ETO treatment. Data represent the mean ± s.e.m. (n = 3 biologically independent samples with more than 600 cells analyzed per treatment). Statistical significance was determined using a two-tailed unpaired Student’s t-test. **P < 0.01; ***P < 0.001.

**Figure S9:**
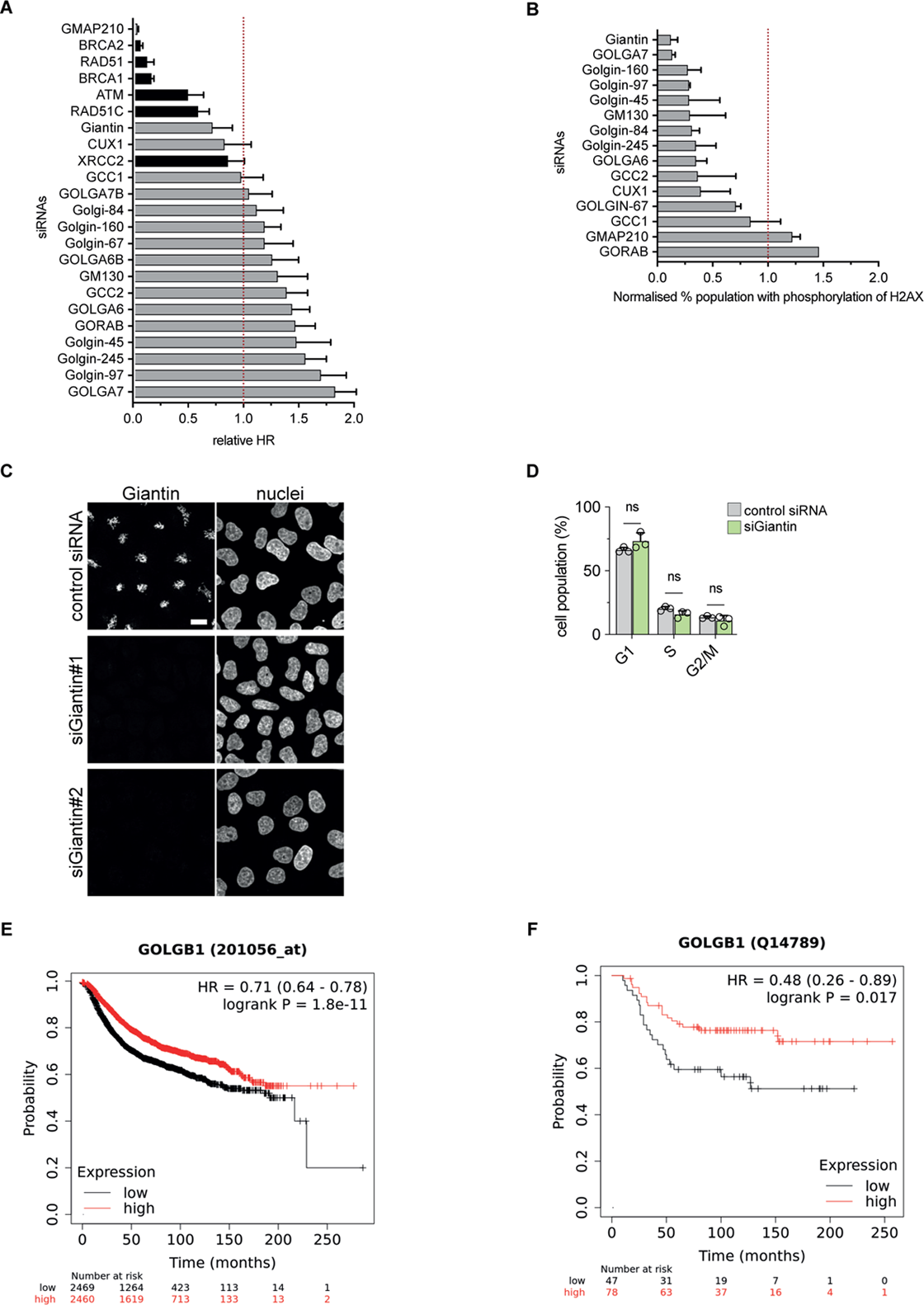
Impact of Golgin proteins on DNA repair, cell cycle, and breast cancer survival, related to Figure 4 and Figure 5. **(A)** Re-analysed siRNA screen data (Adamson et al., 2012) showing the relative Homologous Recombination (HR) repair rate upon systematic knockdown of the Golgin protein family (grey) and HR complex proteins (black). **(B)** Re-analysed siRNA screen data (Paulsen et al., 2009) showing the relative percent cell population with phosphorylation of H2AX upon systematic knockdown of the Golgins. The datasets are normalised to the negative control set at 1. **(C)** HeLa-K cells were transfected with control, or Giantin siRNAs for 72 hours. The cells were stained with antibodies against Giantin and nuclei were stained with Hoechst 33342. Scale bar, 10 μm. **(D)** Cell cycle profile of HeLa-K cell treated with Giantin or control siRNA. Data represent the mean ± s.e.m. (n = 3 biologically independent samples). **(E and F)** Kaplan-Meier survival plots in patients with high and low mRNA **(F)** (Győrffy, 2021) and protein **(G)** (Ősz et al., 2021) Giantin expression in Breast Cancer patients. The graph was generated using the kmplot database (https://kmplot.com).

**Supplementary Table S1.** Set of tables listing the antibodies and siRNA used for the analysis of Human Protein Atlas dataset for the identification and validation of dual-localising Golgi-nuclear proteins. Table 1 presents the antibodies and siRNA used; Table 2 presents the identified and validated Golgi-nuclear protein. Table 3 presents the gene ontology and enrichment terms of the validated Golgi-nuclear proteins.

**Supplementary Table S2.** Table listing the cross-correlation measure for analysis of Golgi cisternal localisation analysis of dual-localising DDR proteins.

